# Oxidative Stress-Responsive Cell Wall Remodeling Depends on Phosphate in *Candida albicans*

**DOI:** 10.64898/2025.12.23.696271

**Authors:** Anand Jacob, Wanjun Qi, Jayasubba Reddy Yarava, Murtaza Barkarar, Aswath Karai, Josh V. Vermaas, Julia R. Köhler, Tuo Wang

## Abstract

The growing number of patients susceptible to invasive *Candida albicans* infections has intensified the need for new antifungal targets in pathways essential for fungal growth and pathogenesis. Among these pathways, phosphate homeostasis has emerged as a significant determinant of virulence, yet how phosphate availability shapes cell wall structure in response to host-derived oxidative stress remains unclear. During commensal growth, *C. albicans* cells typically enjoy phosphate repletion and a neutral redox environment. Transitioning to invade host tissues, they simultaneously experience phosphate deprivation and intense extrinsic oxidative stress. Here, we employ solid-state NMR to render details of cell wall remodeling in response to oxidative stress, in its dependence on phosphate. Phosphate deprived cells remodel the rigid wall core and reduce hydration and polymer mobility in the absence of oxidative stress. During hydrogen peroxide exposure, highly mobile outer polysaccharides are primary interactors. In wildtype cells, some of these polymers are recruited into the rigid core, reinforcing the wall scaffold, whereas phosphate transport mutants fail to undergo this remodeling. These findings establish phosphate acquisition as a component of oxidative defense and link nutrient sensing and -availability to the mechanical resilience of the fungal cell wall, revealing an architectural vulnerability with relevance for antifungal development.

## INTRODUCTION

An important worldwide challenge is the growing number of immunocompromised individuals at risk for invasive candidiasis, driven by therapeutic progress in oncology, rheumatology, gastroenterology, neonatology, and transplantation medicine^1–3^. Despite the increasing clinical need, the current antifungal armamentarium, comprising echinocandins, polyenes, azoles, and the pyrimidine analog 5-flucytosine, is constrained by substantial limitations, including toxicity, pharmacologic interactions, inadequate tissue penetration, and formulations restricted to intravenous or enteral delivery^4,5^. Given the wide range of comorbidities and clinical conditions encountered among patients with invasive candidiasis, no single antifungal agent is suitable for all patient profiles^6^. Compounding these challenges, *C. albicans* strains and other *Candida* species resistant to antifungal agents, particularly azoles and echinocandins, emerge during prolonged treatment courses, and multidrug-resistant strains are associated with increased mortality and healthcare costs^4,5,7–10^. Therefore, *C. albicans* has been categorized as a critical-priority fungal pathogen on the WHO fungal priority pathogens list^11,12^. Given the limited efficacy of existing antifungal therapies in complex clinical scenarios, identification and characterization of new therapeutic targets is urgently needed.

To identify new opportunities for antifungal intervention, it is essential to focus on pathways indispensable for fungal growth and pathogenesis within the host environment, among which phosphate metabolism is particularly compelling. Phosphate is an essential macronutrient for all living organisms, supporting the synthesis of nucleic acids, phospholipids, and ATP, as well as functioning as a key regulator within numerous cellular signaling pathways^13,14^. At the same time, fungal phosphate homeostasis differs fundamentally from that of humans^15^.

*C. albicans* colonizes the gastrointestinal tract of an estimated 50-83% of adults, where it coexists with hundreds of bacterial species in the colon^16,17^. Commensal physiology of *C. albicans* is a distinct state with apparent specific characteristics like suppression of hyphal growth^18,19^. During episodes of disruption of the intestinal epithelial barrier and insufficient response of host neutrophils, the fungus switches its physiology from commensal to infectious, invading host tissues and bloodstream^20^. This transition exposes *C. albicans* to a dramatic change of external conditions, including increased oxygen tension and pH, as well as altered nutrient availability, including phosphate^21^. In addition, invading *C. albicans* cells are typically confronted by host innate immune cells, neutrophils and macrophages.

Human stool contains an average of 11 and up to 33 mM soluble phosphate, indicating that as an intestinal commensal, *C. albicans* experiences phosphate repletion^22^. In contrast, as it invades the tissues and bloodstream of the human host, the fungus experiences phosphate starvation: inorganic phosphate concentration of plasma (which comprises 85% of total plasma phosphate, with the other 15% consisting of organic compounds like glycerophosphocholine) is in the 0.8-1.6 mM range^23,24^. Additionally limiting access to host phosphate is the fact that in the bloodstream and in other extracellular fluid, 15% of inorganic phosphate is protein-bound and another fraction is insolubly complexed to cations like calcium and magnesium. Human body fluids and tissues are slightly alkaline at pH 7.35-7.45, further reducing phosphate accessibility for *C. albicans* whose principal inorganic phosphate transporter, Pho84, has an acidic optimum, a dilemma *C. albicans* shares with other fungal pathogens^25,26^. For these reasons, transcriptional phosphate starvation responses are induced during *C. albicans’* invasive growth in the host^27–29^.

*C. albicans* cells undergoing the transition from commensal to invasive physiology therefore must adapt from a phosphate replete to a phosphate depleted nutritional state. Among genes induced during phosphate deprivation, transcriptional studies have identified cell wall component and remodeling genes, including *MNN1, XOG1,* and *MNN22*^30^. These genes encode an α-1,3-mannosyltransferase, an exo-1,3-β-glucanase and an α-1,2-mannosyltransferase, respectively, and are 18-, 16- and 15-fold upregulated when wildtype cells lack phosphate^30^. β-mannosyltransferases *BMT3* and *BMT4* are 11- and 9-fold upregulated under these conditions, while chitinase *CHT1* and another putative β-mannosyltransferase, *RHD1*, are 4-fold upregulated. Strikingly, putative sensors of cell wall stress are also upregulated, specifically *PGA28* (24-fold up) and *WSC4* (3-fold up), in addition to the marked upregulation of remodeling enzymes, hinting that phosphate scarcity-induced cell wall alterations could predispose to cell wall stress. In fact, cells that lack the major inorganic phosphate transporter Pho84 are hypersensitive to cell wall stress^31^. In addition to Pho84, *C. albicans* encodes three other phosphate transporters that contribute to phosphate acquisition but have more limited physiological roles^25^.

During host tissue and bloodstream invasion, simultaneously with phosphate deprivation and the increased oxygen tension of normally oxygenated blood and tissues, *C. albicans* also encounters intense oxidative stress generated by innate immune cells, neutrophils and macrophages. During the respiratory burst, these phagocytes produce superoxide that is dismutated to hydrogen peroxide, H_2_O_2,_ by enzymes embedded in the cell wall on the fungal surface and in the cytoplasm^32,33^. *C. albicans* deploys an integrated network of antioxidant defenses, including superoxide dismutases, catalase, and the thioredoxin and glutathione systems, which together detoxify ROS and limit cellular damage^34–36^. The activity of these systems is coordinated by stress response signaling pathways such as the Hog1 MAPK pathway and the Cap1 transcriptional regulator^37–41^. In addition, the cell activates protein quality control mechanisms involving the proteasome and initiates DNA repair processes to maintain essential cellular function^42–44^.

Since the wall is the first organelle of the fungal cell to encounter host-derived reactive oxygen species (ROS), it must undergo stress-induced remodeling to maintain structural integrity and modulate immune recognition during infection^45–48^. The cell walls of *Candida* species are organized as a multilayered assembly in which an inner framework of chitin, β-1,3-glucan, and β-1,6-glucan supports an outer layer of mannoproteins extensively decorated with *N*- and *O*-linked mannans^49–52^. Given that phosphate acquisition and signaling via Pho84 are linked to oxidative stress responses, phosphate homeostasis is expected to influence both cell-wall organization and the fungal capacity to withstand host-derived oxidative attack^53^. Most studies probing oxidative stress in *C. albicans* have used high concentrations of H_2_O_2_, which due to their global inhibitory effect do not fully capture active molecular remodeling of cell-wall structures^54–57^. In addition, the phosphate deprivation *C. albicans* experiences in the host simultaneously with oxidative stress, may alter its responses to the latter. We set out to more clearly define the effect of simultaneous oxidative stress and phosphate scarcity on cell wall remodeling, since previous investigations of oxidative stress were performed in the high phosphate conditions of laboratory culture media.

To address this gap, we employed a moderate, sublethal concentration of 0.8 mM H_2_O_2_ to examine oxidative stress responses under physiologically relevant conditions, comparing phosphate-replete with phosphate-deprived cells. Using solid-state NMR, we track structural changes in intact living cells, focusing on the organization and dynamics of glucans and mannans. By comparing the wildtype strain with a mutant lacking all four inorganic phosphate transporters, we are able to directly assess how phosphate availability shapes oxidative stress responses and cell-wall remodeling. This strategy enables the identification of phosphate-dependent mechanisms that support fungal survival during immune challenge and may reveal structural vulnerabilities suitable for antifungal intervention.

Such efforts are made possible by recent advances in solid-state NMR methodologies, which now allow in situ analysis of cell-wall composition, hydration, and dynamics across a broad range of fungal species^58–60^. Recently, these approaches have been applied to *Aspergillus* (*A. fumigatus*, *A. nidulans*, and *A. sydowii*)^61–67^, other filamentous fungi including *Neurospora*, *Mucor*, *Rhizopus*, and *Schizophyllum*^68–71^, and yeast cells of *Cryptococcus*^72–75^, *Schizosaccharomyces*^76^, and *Candida* species^77^. In particular, recent in situ solid-state NMR work showed that although *C. albicans* and *C. auris* share the same hierarchical cell-wall architecture, their remodeling strategies diverge under echinocandin exposure: *C. albicans* thickens and reinforces its rigid core, whereas *C. auris* maintains wall integrity by upregulating β-1,6-glucan biosynthesis^77^. These findings provide a structural framework for interpreting stress-dependent remodeling in *Candida* cell walls, including the responses to ROS and phosphate deprivation evaluated in this work.

Building on this foundation, here we use multidimensional ^13^C and ^1^H-detected solid-state NMR to directly observe how severely restricted phosphate uptake alters cell-wall structure in intact cells of *C. albicans*. Although Pho84 plays the dominant role in all phosphate availability-linked processes^25^, we use a mutant lacking all four inorganic phosphate transporters mutant (*pho84Δ/Δ pho89Δ/Δ pho87Δ/Δ fgr2Δ/Δ*) to maximize the structural phenotype detectable by NMR analysis. Our results demonstrate that phosphate availability reshapes cell-wall architecture and modulates the oxidative-stress response, revealing a mechanistic link between phosphate metabolism and cell wall resilience.

## RESULTS

### Phosphate transport deficiency fully remodels the rigid cell-wall core in *C. albicans*

The ^13^C-labeled cells of *C. albicans* were kept intact and viable during solid-state NMR analysis. Prior to isotopic labeling, the cultures were subjected to sequential phosphate-deprivation cycles to establish a defined phosphate-starved physiological state, ensuring that the observed cell wall features arise from altered phosphate acquisition and signaling rather than from remaining intracellular or extracellular phosphate reserves. Without chemical perturbations, all molecules preserved their native properties and supramolecular organization. Using a variety of polarization techniques, we can achieve selective detection of molecules across distinct dynamical regimes (**Supplementary Fig. 1**). The rigid fraction was selectively detected through ^1^H-^13^C cross polarization (CP), which favors rigid molecules exhibiting large C-H dipolar couplings, and was projected into a 2D ^13^C-^13^C correlation spectrum for improved spectral resolution. In wildtype cells, this method predominantly detected β-1,3-glucan signals (**Fig. 1A**). Representative cross peaks include correlations of carbon 1 with carbons 5, 2, and 4 (B1-5, B1-2, and B1-4), and correlations involving carbon 6 (B4-6, B5-6, and B1-6), enabling assignment of ^13^C chemical shifts for individual carbon sites within the β-1,3-glucan structure (**Fig. 1B**).

**Figure 1.**
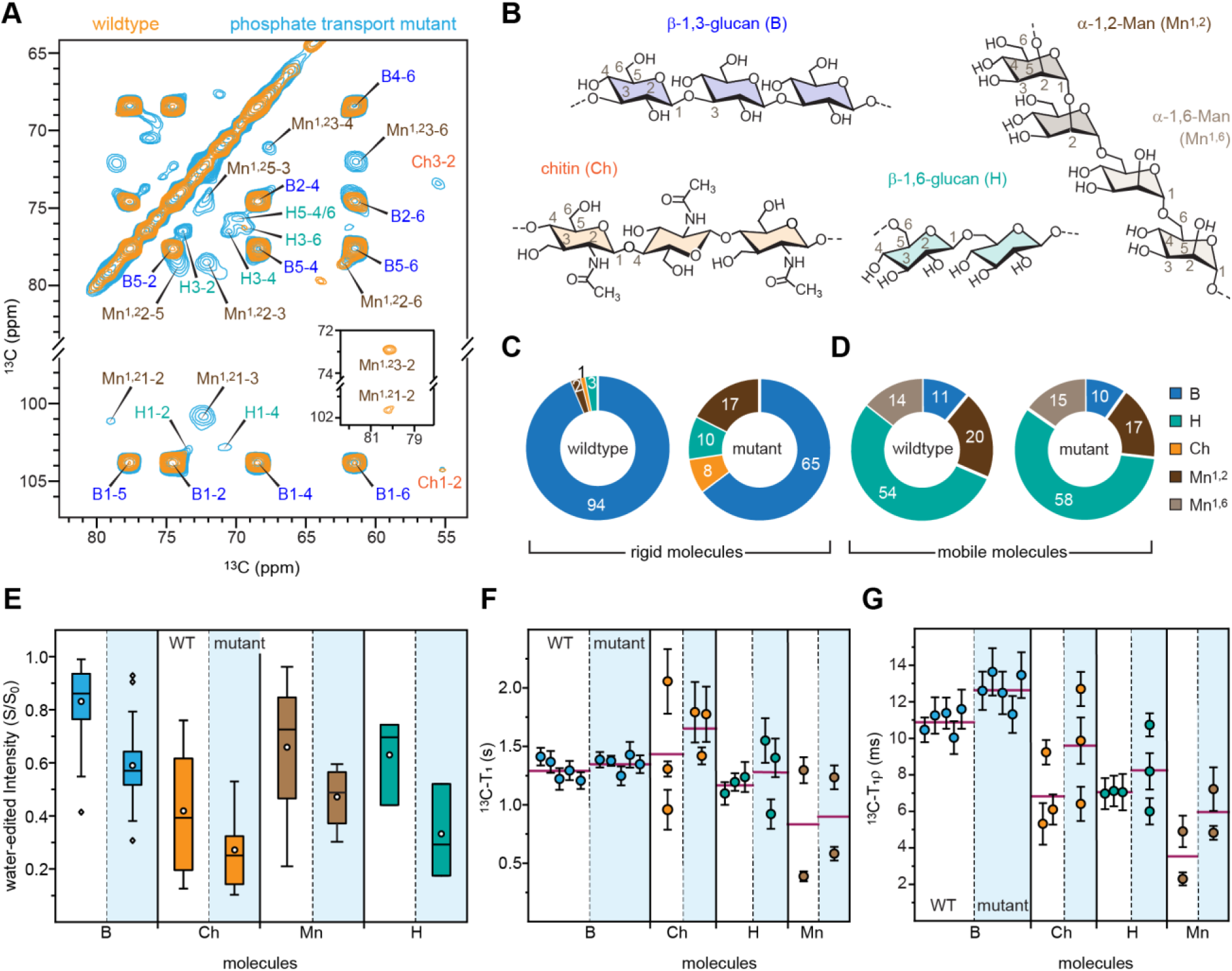
Phosphate-transport mutant of *C. albicans* displays altered cell wall structure. (**A**) Rigid-polysaccharide carbon signals detected by 2D ^13^C-^13^C CORD spectra from wildtype (JKC 915; orange) and phosphate transport mutant (JKC 2830; light blue) *C. albicans* cell walls. The inset shows weak mannose signals in the wild-type sample, which are only visible after lowering the contour levels of plotting. NMR abbreviations are used for resonance assignment. β-1,3-glucan: B; chitin: Ch; β-1,6-glucan: H; α-1,2-mannose units: Mn^1,2^; α-1,6-mannose units: Mn^1,6^. For example, B1-2 denotes the cross peak between carbons 1 and 2 in β-1,3-glucan. (**B**) Simplified structures of major polysaccharides. Molar compositions of (**C**) rigid polysaccharides and (**D**) mobile carbohydrates were estimated from resolved cross-peak volumes in 2D ^13^C CORD spectra and DP-refocused J-INADEQUATE spectra, respectively. Numbers represent percentages. (**E**) Box-and-whisker plot showing the water-edited intensities (S/S_0_) of β-1,3-glucan (blue), chitin (orange), mannan (brown), and β-1,6-glucan (green) in wildtype cells (WT; white background) and in the phosphate-transport mutant (light-blue background). For each sample, n=36 for β-1,3-glucan, n=5 for chitin, n=4 for mannan, and n=3 for β-1,6-glucan. The boxes represent the IQR, with whiskers extending to 1.5 IQR. For β-1,6-glucan, all data points are within the range between the first and third quartile, thus there are no whiskers. Mean values are represented by open circles and medians by purple horizontal lines. (**F**) ^13^C-T_1_ and (**G**)^1^H-T_1ρ_ relaxation time constants of rigid cell wall polysaccharides in wildtype cells (white background) and in the phosphate-transport mutant (light-blue background). For both panels, error bars represent s.d. of the fitted parameters, and the horizontal lines in magenta indicate the average relaxation time constant for each polymer. Source Data are provided as a source data file.

Surprisingly, the rigid fraction of the phosphate transport mutant exhibited substantially increased spectral complexity. In addition to strong β-1,3-glucan signals, the chitin (*e.g.*, Ch3-2 and Ch1-2) and β-1,6-glucan (*e.g.*, H3-6) signals were significantly more intense than in the wildtype. Moreover, signals corresponding to α-1,2-linked mannose units, which form diverse sidechains decorating the α-1,2-linked backbone of mannan in *C. albicans* (**Fig. 1B**), became prominent in the mutant. Notable examples include cross peaks between carbons 1 and 3 (Mn^1,21-3^) and between carbons 2 and 3 (Mn^1,22-3^) (**Fig. 1A**). Peak-intensity analysis showed that mannan, chitin, and β-1,6-glucan constitute 17%, 8%, and 10% of the rigid polymers in the mutant, respectively, whereas the wildtype cell wall is dominated by β-1,3-glucan (94%) and contains only small amounts of chitin (1%), β-1,6-glucan (3%), and mannan sidechains (2%) in the rigid fraction (**Fig. 1C**).

In contrast, no substantial changes were observed in the mobile fraction. This fraction was selectively detected using a combination of ^13^C direct polarization (DP) and a short recycle delay, which preferentially excites molecules with rapid ^13^C-T_1_ relaxation and suppresses signals from rigid components that relax slowly. This polarization scheme, combined with 2D refocused J-INADEQUATE spectroscopy, enabled unambiguous identification of the carbon signals and connectivities in mobile cell-wall polymers (**Supplementary Fig. 2**). We identified both the α-1,2-linked backbone and α-1,6-linked mannan sidechains, as well as β-1,6-linked and β-1,3-linked glucans and their β-1,3,6-linked branching sites. Intensity analysis revealed similar mobile-domain compositions for both wildtype and mutant cells: 54-58% β-1,6-glucan, 32-34% mannan, and 10-11% β-1,3-glucan (**Fig. 1D**).

Because the rigid fraction plays a central role in cell-wall remodeling in the phosphate transport mutant, we next examined how the phosphate transport defect perturbs hydration and dynamics of rigid biopolymers. Hydration profiles were assessed using water-to-carbohydrate ^1^H polarization transfer experiments, where the S/S_0_ ratio (water-accessed versus control) reflects water accessibility at specific carbon sites (**Supplementary Fig. 3**). These ratios were considerably lower in the mutant than in the wildtype, decreasing from 0.83 to 0.64 for β-1,3-glucan and from 0.41 to 0.27 for chitin (**Fig. 1E**). The few weakly resolved peaks attributed to β-1,6-glucan and mannan sidechains exhibit higher S/S_0_ ratios than chitin but lower than those of β-1,3-glucan. Moreover, both β-1,6-glucan and mannan show a decrease in the S/S_0_ ratio. These observations are reasonable because β-1,3-glucan has recently been identified as a key determinant of cell-wall hydration across diverse fungal species. Therefore, the one-third reduction of β-1,3-glucan in the mutant cell wall inevitably compromises the ability of carbohydrate polymers in the rigid core of the *C. albicans* cell wall to retain water molecules.

The motions of polysaccharides were probed on two distinct timescales: ^13^C-T_1_ relaxation to assess local nanosecond dynamics and ^1^H-T_1ρ_ relaxation to probe larger-scale collective motions on the microsecond timescale (**Supplementary Fig. 4**). Chitin showed marked increases in the averaged values of both ^13^C-T_1_—from 1.6 s in the wildtype to 2.1 s in the mutant (**Fig. 1F**)—and ^1^H-T_1ρ_—from 6.8 ms to 9.6 ms (**Fig. 1G**), signifying reduced mobility on both timescales. In contrast, β-1,3-glucan only exhibited an increase in ^1^H-T_1ρ_ from 10.8 ms to 12.6 ms, indicating enhanced microsecond-scale dynamics without corresponding nanosecond -scale restriction.

The small amount of mannan side chains detectable in the rigid phase remains relatively mobile, as evidenced by very short ^13^C-T_1_ and ^1^H-T_1ρ_ relaxation time constants. This indicates that partial mobility persists even after these sidechains come into close proximity with—and become partially rigidified by—chitin and β-1,3-glucan. In contrast, β-1,6-glucan exhibits ^13^C-T_1_ values comparable to those of β-1,3-glucan but shows greater nanosecond -scale dynamics, as reflected by a shorter ^1^H-T_1ρ_. Both molecules also showed an increase in relaxation time constants from the wildtype to the mutant.

In summary, the phosphate transport defect profoundly remodeled the rigid core of the *C. albicans* cell wall. This remodeling involves reshaping both the molecular composition and the physical properties of the rigid scaffold. Whereas the wildtype rigid cell wall is dominated by β-1,3-glucan, the mutant incorporates increased levels of chitin and β-1,6-glucan, together with some mannan sidechains that interact with these components. The cell wall also exhibits decreased water accessibility and reduced molecular mobility, resulting in a stiffer, less permeable barrier. This remodeling strategy in response to phosphate deprivation resembles cell-wall adaptations triggered by external stressors, such as antifungal treatment with echinocandins or exposure to a hypersaline environment^61,64,70,77^.

### 1H-detected solid-state NMR reveals reduced polymorphism in mobile polysaccharides

It is fully unexpected that molecules in the mobile fraction did not exhibit substantial changes in composition. One possibility is that more subtle structural variations, such as conformational changes, occurred but escaped detection in ^13^C-based experiments due to limited spectral resolution. Therefore, we applied ^1^H-detected solid-state NMR spectroscopy to take advantage of its enhanced spectral sensitivity and the strong dependence of ^1^H chemical shifts on subtle carbohydrate structural differences. This approach enables improved resolution of the polymorphic forms of mobile polysaccharides, which are often poorly distinguished in ^13^C-detected spectra due to extensive chemical-shift degeneracy.

The 2D J-HCCH TOCSY spectra of intact *C. albicans* cells revealed marked differences in the carbohydrate anomeric region between wildtype and mutant strains (**Fig. 2A**). In particular, the α-1,2-linked mannose units showed a substantial reduction in peak multiplicity in the phosphate transport mutant. For example, the Mn^1,2^1^m^ and Mn^1,2^1^j^ peaks corresponding to the C1/H1 sites of the type-m and type-j forms of α-1,2-linked mannose residues completely disappeared. In addition, signals corresponding to the type-j, type-k, and type-i forms merged into a single continuous band in the mutant, rather than three discrete peaks as observed in the wildtype cells. Spectral simplification was also observed for the Gl1 signals corresponding to C1/H1 of Glc/Gal and their derivatives, where type-f and type-b were absent in the mutant. β-1,3-glucan also exhibited reduced structural complexity in the mutant, as demonstrated by the decreased number of peaks for its C3/H3 sites. Signals corresponding to type-d units and branching sites were weaker or absent in the mutant. These results indicate that phosphate transport deficiency leads to a more homogeneous molecular structure within the mobile matrix of the cell wall, despite the overall molecular composition remaining unchanged. This suggests that either the modification of mobile polysaccharides is impeded, reducing their chemical diversity, or that the polymers adopt more homogeneous conformations due to simplified intermolecular interactions and enrichment of energetically favored conformations.

**Figure 2.**
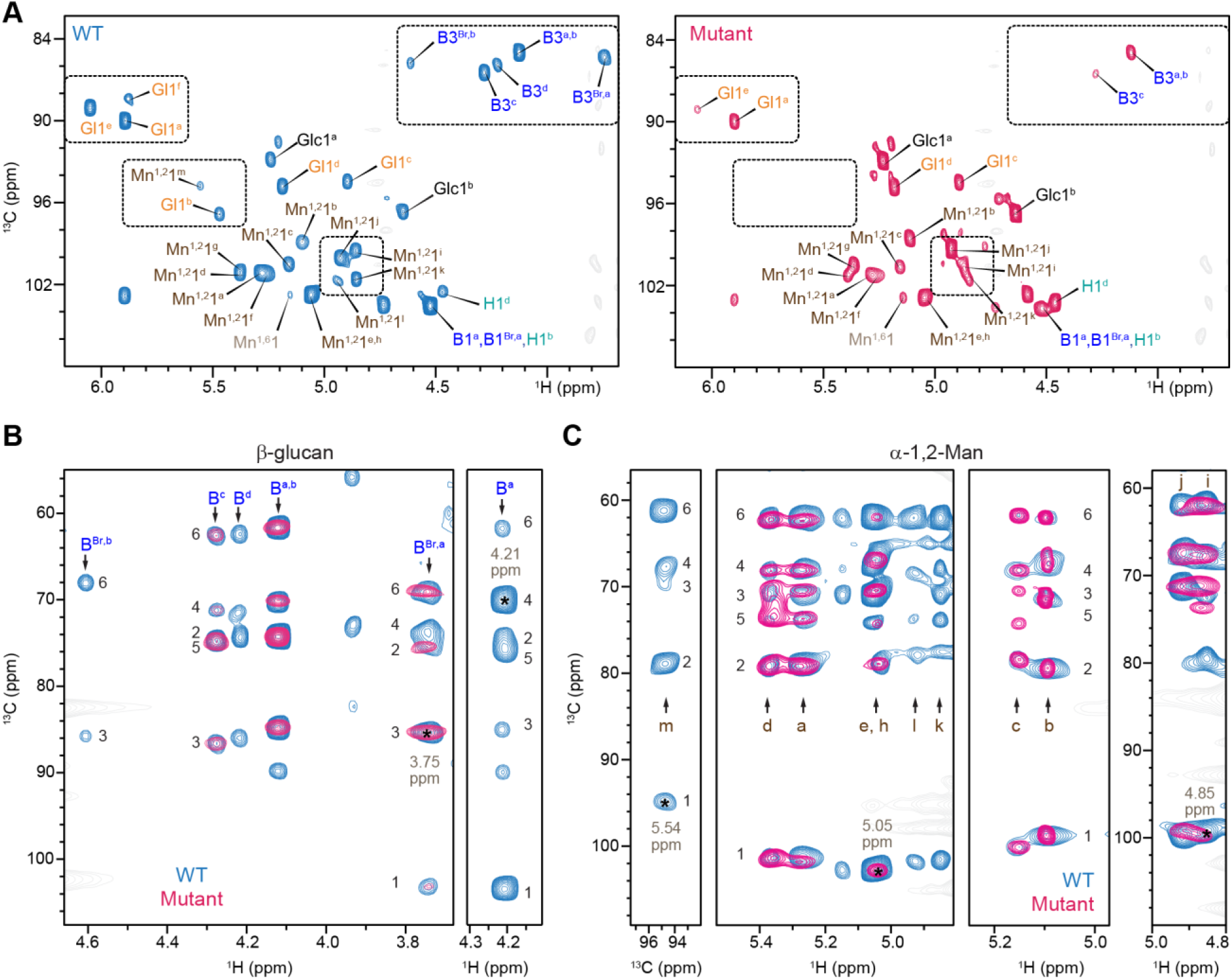
Proton-detected solid-state NMR shows reduced structural polymorphism in mutant walls. (**A**) Comparison of 2D hcCH TOCSY (DIPSI-3) spectra of wildtype (blue, left) and phosphate transport mutant (magenta, right) strains of *C. albicans*, showing one-bond ^13^C-^1^H correlations. Water signals are in grey. Dotted boxes highlight spectral regionswherethe mutant displayed fewer carbohydrate peaks relative to wildtype cells. Mn: mannan; B: β-1,3-glucan; B^Br^: branching site of the 1,3,6-Glc unit; H: β-1,6-glucan; Mn1,2: α-1,2-Man; Mn1,6: α-1,6-Man; Glc: glucose; Gl: galactose or glucose derivatives. Numbers indicate the carbon/proton position, and superscript letters denote different structural forms of a molecule (*e.g.*, B3ᵈ denotesthe C3-H3 cross peak belonging to type-d β-1,3-glucan). (**B**) β-glucan and (**C**) α-1,2-Man exhibit multiple structural allomorphs as resolved from ^13^C-^1^H and ^13^C-^13^C strips extracted from 3D hCCH TOCSY (DIPSI-3) spectra. The spectra show through-bond carbon-carbon connectivity (carbon numbers labeled) for each molecule or structural formalong the vertical signal traces. The strip extracted at a specific carbon or proton site is indicated by an asterisk. In wildtype cells (blue), resolved signals are assigned to three forms of β-1,3-glucan (B^a^, B^b^, and B^c^) and two forms of branching sites in β-1,3-glucan (B^Br,a^ and B^Br,b^) in Panel B, and eleven forms of α-1,2-Man units with well-resolved carbon connectivity in Panel C. The phosphate-transport mutant (magenta) displayed substantially reduced structural polymorphism, with only two types of β-1,3-glucan, one form of the branching site, and seven of the α-1,2-Man forms detected.

Extending the 2D experiment into its 3D format, the J-HCCH TOCSY experiment incorporating DIPSI-3 mixing enabled tracking of carbon connectivities through J-coupling and allowed unambiguous site-specific assignments for each allomorph^65,75,78^. For β-1,3-glucans, type-a and type-b displayed overlapping signals (B^a,b^) in both samples in the stripe extracted at ^1^H chemical shift of 3.75 ppm. However, the characteristic type-a peaks (B^a^) extracted at ^1^H chemical shift of 4.21 ppm was only detected in the wildtype, indicating that type-b is present in both strains whereas type-a is confined to the wildtype. Type-c (B^c^) was detectable in both strains but was weaker in the mutant, and type-d (B^d^) was completely absent in the mutant, consistent with the 2D spectra. The first type of branching site B^Br,a^ was observed in both strains, although weaker in the mutant, while the second type B^Br,b^ was not detected in the mutant. These findings revealed a simplified β-1,3-glucan matrix, with reduced complexity in both the linear chains and the branching patterns, forming a more uniform structure across the *C. albicans* cell wall.

As mannan fibrils form the outer layer of the *C. albicans* cell wall, their sidechains, predominantly comprising α-1,2-linked mannose units, play a critical role in stabilizing this layer through extensive intermolecular interactions. This leads to remarkable structural diversity of these sidechains, reflected in the coexistence of 14 resolvable allomorphs in the spectra. Representative stripes from the 3D J-HCCH TOCSY spectra are shown in **Fig. 2C**, where the mutant has completely lost the signals corresponding to forms m, l, and k, and the j and i forms are substantially restructured, consistent with the 2D results but now with improved resolution and complete resonance assignments.

Together, these findings demonstrate that the phosphate transport defect significantly reduces the structural complexity accommodated by the mobile domain of the *C. albicans* cell wall, even without major perturbations to molecular composition. They also highlight the power of ^1^H-detected solid-state NMR for resolving polysaccharide polymorphism and subtle structural modifications at high sensitivity and resolution in intact cells.

### Oxidative stress induces distinct changes to polysaccharides in *C. albicans* cell walls

To assess the impact of oxidative stress on *C. albicans* cell wall architecture, both wildtype and phosphate transport deficient mutant cells were treated with 0.8 mM H_2_O_2_, a concentration selected to permit sustained growth of treated cells. Sublethal, sub-millimolar H_2_O_2_ elicits adaptive responses in *C. albicans* and may reflect the ROS levels generated during neutrophil and macrophage respiratory bursts in the host, enabling biologically meaningful oxidative stress without catastrophic loss of viability^36,55,79–82^. The aim of this experiment was to isolate oxidative stress-specific remodeling of the cell wall during growth of phosphate deprived cells. For this reason, the phosphate transport defective mutant and wildtype cells were cultured in standard, phosphate replete media for 8 hours. In this medium, mutant cells are able to import some inorganic phosphate via their glycerophosphocholine transporters^25^. Comparative analyses revealed marked differences in polysaccharide behavior that depend on polymer dynamics (**Supplementary Figs. 5** and **6**) and, likely, cell-wall organization. Surprisingly, we also observed perturbations that can only have arisen from reaction of peroxide directly with cell wall components. This was despite the fact that due to its expected spontaneous disproportionation in the watery medium, no H_2_O_2_ was detectable in the medium at the time of the cell harvest. Hence we attribute the observed cell wall alterations to two factors, firstly, active remodeling of the wall by cells responding to oxidative stress, and secondly chemical reaction of peroxide with cell wall components.

Highly mobile and largely solvated carbohydrates, detected using the 1D refocused INEPT experiment (**Fig. 3A**), and the overall mobile fraction, detected using 1D ^13^C DP spectra (**Fig. 3B**), showed the characteristic signals of α-1,2-Man and α-1,6-Man in mannan sidechains and backbones, β-1,6-glucan, and partially β-1,3-glucan in the wildtype cells. This agrees with their localization: mannan is surface-exposed, and β-glucans are dynamically flexible and highly hydrated. After H_2_O_2_ exposure, these signals completely disappeared in both wildtype and mutant cells in the highly mobile fraction detected by INEPT (**Fig. 3A**). This is a signature feature of paramagnetic relaxation enhancement (PRE)^83,84^, in which NMR peaks are severely broadened by rapid transverse relaxation arising from the presence of paramagnetic species, such as radicals formed when carbohydrates react with H_2_O_2_, which generates hydroxyl and related radicals capable of abstracting hydrogen from polysaccharide residues, forming carbon or oxygen centered radicals^85–89^. Peroxide-induced structural changes to polysccharides such as hydrogen abstraction, degradation, or cross-linking may further contribute to the signal loss, but to a lesser extent compared to paramagnetic relaxation enhancement. The remaining signals were dominated by lipids, which are located deeper in the fungal cell membrane below the cell-wall layers.

**Figure 3.**
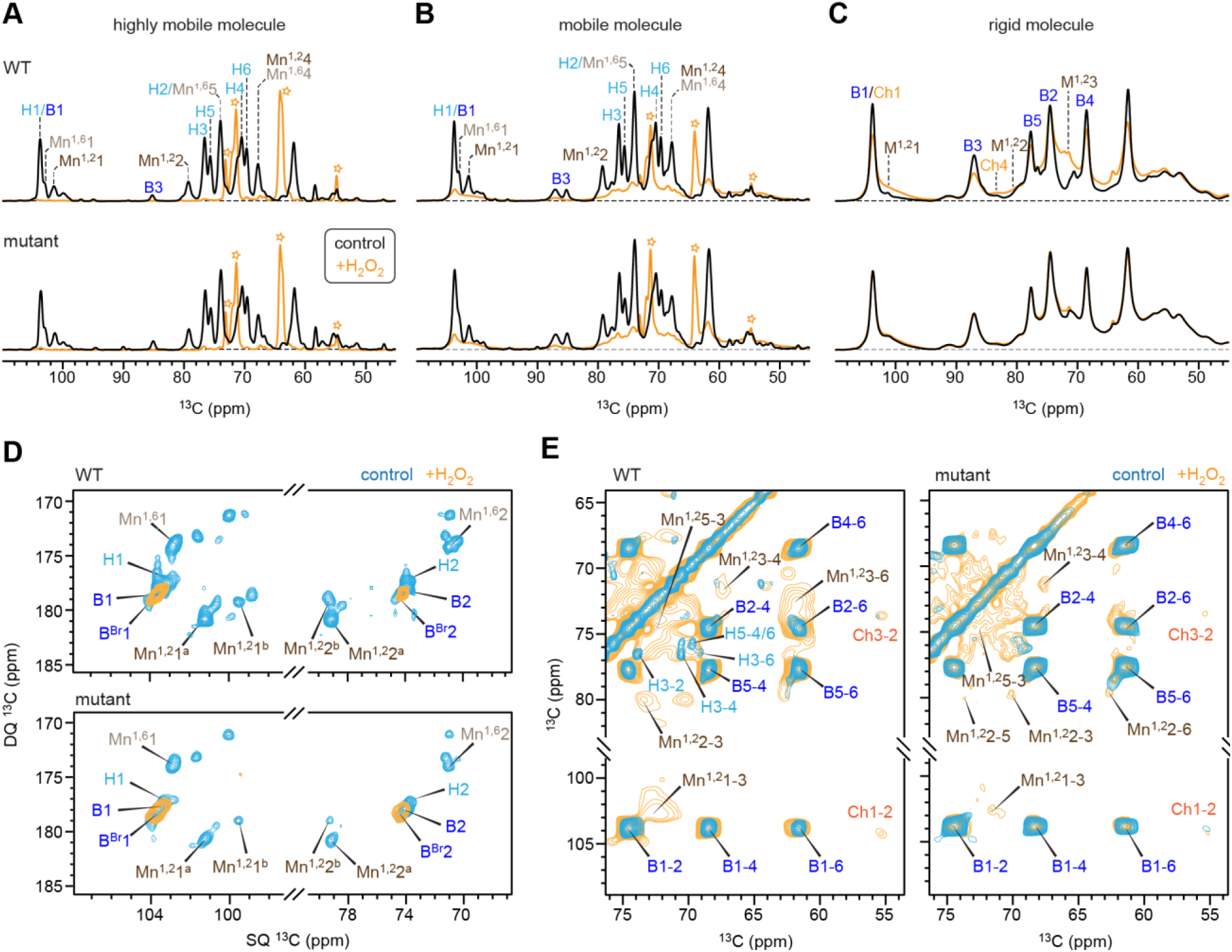
Alterations of polysaccharides in C. albicans cell walls during oxidative stress. 1D ^13^C NMR spectra were obtained using (**A**) INEPT to detect highly mobile polysaccharides, (**B**) DP to detect mobile polysaccharides, and (**C**) CP to detect rigid polysaccharides. Spectra are shown for wildtype (top) and mutant (bottom) cells under control (black) or H_2_O_2_-treated (orange) conditions. Reaction of H_2_O_2_ with carbohydrates generates paramagnetic species (free radicals), which broaden out the signals of associated carbohydrates. Dashed horizontal lines denote the spectral baseline. Open stars mark predominant lipid signals in the INEPT and DP spectra of H_2_O_2_-treated samples, where carbohydrate signals are broadened out due to their interactions with H_2_O_2_. (**D**) mobile molecules detected through 2D ^13^C DP refocused J-INADEQUATE spectra of wildtype (top) and mutant (bottom) cells under control (blue) or H _2_O_2_-treated (orange) conditions. H_2_O_2_ treatment nearly eliminates all mobile polysaccharide signals, except for β-1,3-glucans. (**E**) 2D ^13^C-^13^C CORD spectra of wildtype (top) and mutant (bottom) cells under control (blue) or H_2_O_2_-treated (orange) conditions. New mannan signals appear after H_2_O_2_ treatment, indicating a transition from the mobile fraction to the rigid cell-wall core.

Carbohydrates in the overall mobile fraction detected by ^13^C DP combined with short recycle delays also exhibited substantial reduction in signal intensity (**Fig. 3B**), but unlike the highly mobile fraction, these signals were not entirely lost. A residual pattern of broad peaks remained after treatment, indicating the presence of semi-mobile molecules with restricted accessibility to peroxides. These observations were consistent across both wildtype and mutant cells.

In contrast, rigid molecules detected by ^13^C CP showed only moderate line broadening in the wildtype (**Fig. 3C**). A more pronounced change occurred in mannan, where certain α-1,2-Man units displayed stronger signals in the rigid fraction. The phosphate transport mutant, however, exhibited little to no radical-induced paramagnetic broadening among mannan and glucan peaks and showed reduced compositional changes in the rigid fraction, suggesting a decreased remodeling response to oxidative stress.

Higher-resolution analysis of mobile molecules using 2D ^13^C DP-INADEQUATE spectra revealed that signals corresponding to β-1,6-glucan and mannan were completely absent after oxidative stress, whereas β-1,3-glucan signals partially survived (**Fig. 3D** and **Supplementary Fig. 7**). The rigid fraction showed more complex changes: the 2D ^13^C-^13^C CORD spectra revealed strong paramagnetic broadening in the wildtype, particularly for α-1,2-mannose units and β-1,6-glucan (**Fig. 3E**). The severe broadening of α-1,2-mannan resonances in the wildtype indicates direct interaction of peroxide and the radicals generated upon reaction with this polysaccharide. In contrast, the mannan signals remained substantially sharper in the phosphate transport mutant, confirming that mannans in the rigid core of the mutant cell wall did not react with peroxide to generate radicals.

Taken together, these findings provide new insights into how the cell wall is remodeled physiologically, versus during phosphate scarcity, in response to ROS. In the wildtype, the inner core undergoes significant reorganization, in which a portion of the originally mobile α-1,2-mannan and β-1,6-glucan is incorporated into the rigid core where it interacts with chitin and β-1,3-glucan to reinforce the structural scaffold. In the phosphate-transport mutant, peroxide induces much weaker reinforcement of the rigid core. Although α-1,2-mannan and β-1,6-glucan also redistribute partially into the rigid core after H_2_O_2_ exposure, the extent is substantially reduced.

Peroxide preferentially targets the mobile, surface-exposed polysaccharides, primarily mannan, and the underlying β-1,6-glucans in the intermediate layer of the *C. albicans* cell wall. Thus, the flexible outer polysaccharide layer is highly vulnerable to oxidative damage. These results demonstrate that cell wall remodeling during the physiologic oxidative stress response requires phosphate repletion of the cell. Defective phosphate transport impairs the cell wall remodeling processes that optimize *C. albicans’* ability to withstand oxidative stress.

Interestingly, when *C. albicans* cells previously exposed to hydrogen peroxide were profusely washed, the post-wash spectra showed sharp, well-resolved resonances, particularly for dynamic polysaccharides such as α-1,2-mannose and β-1,6-glucan (**Supplementary Figs. 8-10**). This recovery indicates efficient quenching of radicals generated during peroxide exposure and confirms that the peroxide-induced loss of NMR signals was primarily due to paramagnetic relaxation enhancement. These carbohydrate radicals appear to be transient and can be removed by simple washing, restoring hydroxyl groups or causing only minimal modification, while irreversible processes such as covalent cross-linking or bond cleavage have not occurred, or have occurred only in a very small fraction of molecules. These findings demonstrate that oxidative damage to the mobile domain of the cell wall, mediated through radicalization of flexible mannans and branched glucans, is largely reversible.

### Molecular Dynamics simulation reveals non-specific H_2_O_2_ reactivity with polysaccharides

When examining the direct chemical effects of peroxide on cell wall components, the next question was whether the distinct oxidative responses of cell-wall polysaccharides arise from intrinsic chemical reactivity or from different degrees of accessibility within the differently organized cell walls of wildtype and mutant strains. To evaluate whether particular polysaccharides or structural motifs are preferentially exposed to oxidative attack, we performed molecular dynamics (MD) simulations to assess the interactions between hydrogen peroxide and representative fungal polysaccharides in water (**Supplementary Fig. 11**). These included chitin, mannan, β-1,6-glucan, and β-1,3-glucan incorporating branched structures (**Fig. 4A**).

**Figure 4.**
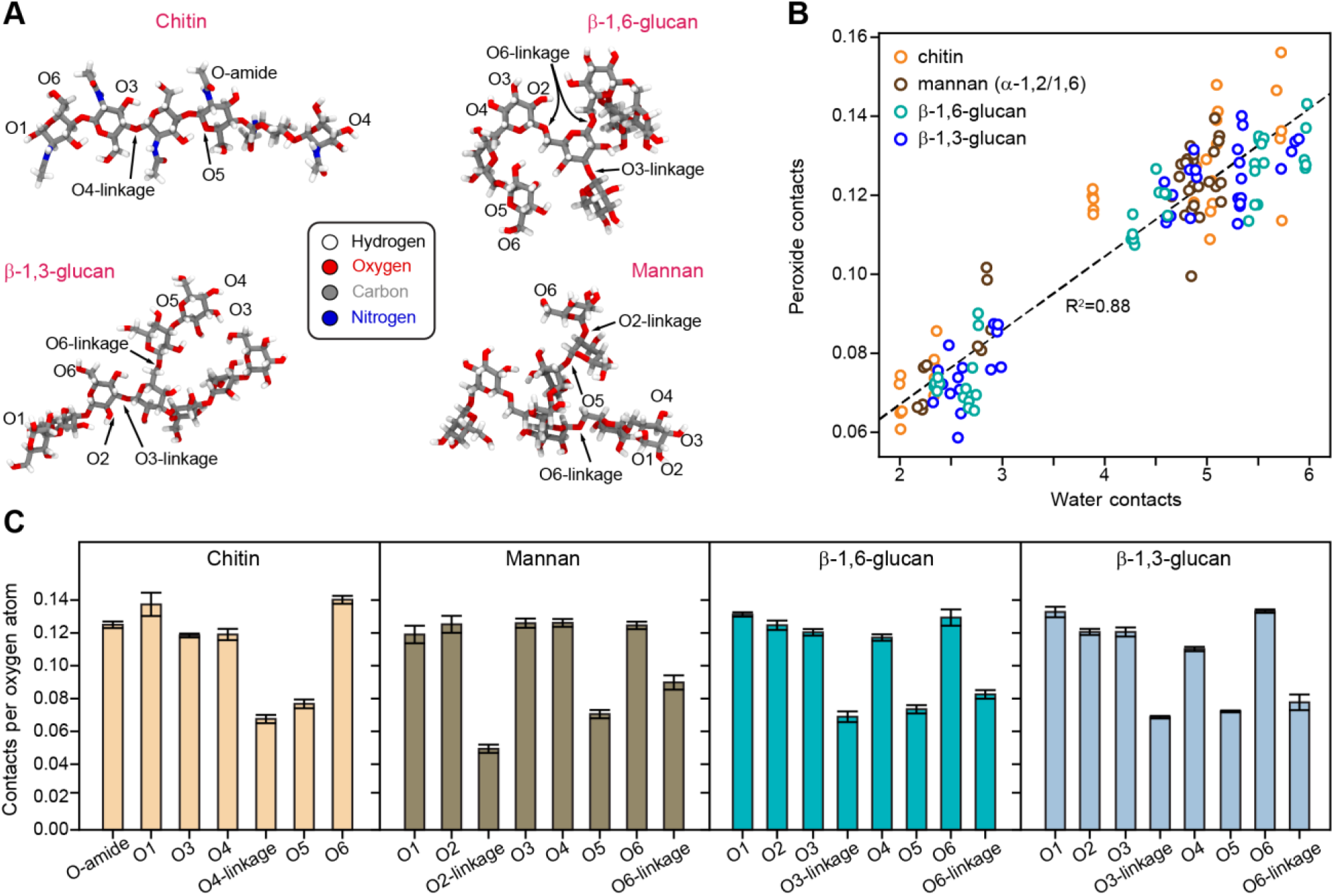
MD simulations reveal peroxide accessibility independent of polysaccharide type. (**A**) Representative MD snapshots illustrating the atom-naming conventions used in this study, highlighting the specific oxygen atoms that can interact with H_2_O_2_. (**B**) Scatter plot comparingthe contacts that each labeled oxygen atom forms with either water or hydrogen peroxide during the simulations. Each point represents a single simulation and specific oxygen position; data points are colored by polysaccharide chain type. Dashed line represents the linear regression, which demonstrates a strong fit with an R^2^ value of 0.88. (**C**) Quantification of H_2_O_2_ contactsper oxygen atom per simulation snapshot across the four glucan chain types analyzed. The error bars represent the standard error, computed based on the variance observed across the five simulation repeats for each glucan chain type.

Across all examined oxygen sites, including those within glycosidic linkages, the frequency of peroxide contacts varied only modestly in the range of 0.06-0.16 (**Fig. 4B**). Even oxygen atoms expected to be sterically shielded, such as those participating in backbone linkages, showed only slightly reduced interactions with peroxide. Consistent with this, a strong linear correlation was observed between the number of water contacts and peroxide contacts. Oxygen sites that were more frequently hydrated were also more frequently contacted by peroxide, indicating that interaction is governed primarily by solvent accessibility rather than inherent chemical preference. Thus, to first approximation, all oxygen atoms within these polysaccharides are susceptible to oxidative attack.

A detailed comparison of individual oxygen positions revealed a consistent trend across all polysaccharides (**Fig. 4C**). Oxygen atoms forming the glycosidic linkages (e.g., O3 and O6 in β-glucans, O2 and O6 in mannan, and the O4-linkage in chitin) exhibited fewer peroxide contacts than the hydroxyl oxygens. O5, which is involved in ring closure and is less solvent exposed, showed a similar decrease. Aside from these predictable reductions, the overall range of peroxide contacts was highly similar among chitin, mannan, β-1,6-glucan, and β-1,3-glucan. This consistency demonstrates that the detailed chemistry of the carbohydrate backbone does not strongly bias peroxide association. Instead, solvent exposure of hydroxyl groups is the dominant determinant of peroxide interaction, regardless of polysaccharide type. Although the absolute differences in contact frequency are small, the extensive sampling in the simulations ensures that the observed trends are statistically robust. Consequently, the most probable sites of glucan peroxidation correspond to the most solvent-accessible oxygen atoms, reinforcing the notion that direct oxidative damage preferentially targets the exposed and dynamic domains of the fungal cell wall.

## DISCUSSION

*Candida albicans* is the most commonly isolated invasive fungal pathogen of humans. This is because, unlike environmental fungal pathogens like the *Aspergilli* and *Mucorales*, it is already present on most humans’ mucous membranes and hence is poised to invade their tissues and bloodstream when epithelial barriers and innate immune defenses weaken. However, the transition from life as a commensal to life as an invasive pathogen exposes *C. albicans* cells to dramatic changes in environmental conditions. In addition to increased ambient oxygen tension and pH, *C. albicans* cells experience sharp nutritional shifts including phosphate scarcity.

Simultaneously with phosphate deprivation during the transition from commensal to invasive pathogen, *C. albicans* in most human hosts experiences the attack of innate immune cells that unleash superoxide anions and antimicrobial peptides onto the fungal cell. Superoxide dismutases at the fungal surface convert this product of host cells’ NADPH oxidase into H_2_O_2_, which diffuses readily across membranes and induces intracellular oxidative stress^90–92^. Given the importance of ROS in the host-pathogen interaction, numerous studies have focused on the impact of oxidative stress on *C. albicans* cells. They have, however, examined oxidative stress responses during the phosphate repletion state that *C. albicans* experiences as a commensal, while the constraints to its oxidative stress responses during the phosphate scarcity typical for its invasive infectious lifestyle have not been investigated.

As a large organelle that interfaces directly with the host, the cell wall is particularly important for *C. albicans*’ ability to withstand stress. Its three wall-anchored superoxide dismutases are a first line of defense against host-generated superoxide^90^. The H_2_O_2_ produced by their catalytic activity may diffuse away, may enter the *C. albicans* cell and cause internal oxidative stress, and may, as we now find, directly alter the chemistry of the structural cell wall polysaccharides.

Intracellular ROS like H_2_O_2_ and the product of its reaction with iron-sulfur clusters, the hydroxyl anion, inflict oxidative damage on membrane lipids, nucleic acids and proteins. Signaling pathways mediated by the stress-activated kinase Hog1 and the transcription factor Cap1 coordinate induction of protective genes: for example the *hog1*Δ mutant is hypersensitive to ROS^39^. Additional regulators such as the zinc-finger factor Sfp1 and the DNA-binding protein Rap1 have been implicated in fine-tuning ROS response and antioxidant gene expression, linking oxidative stress defense to global transcriptional and metabolic regulation^93^. In this way, protective intracellular enzymes are induced, like superoxide dismutases, catalase, glutathione- and thioredoxin-dependent systems, which detoxify ROS and repair oxidative damage^94^. Overall, ROS remain a potent challenge to *C. albicans*: combined stresses (*e.g.*, oxidative plus cationic stress) can suppress antioxidant responses (via inhibition of Cap1 nuclear accumulation) and kill fungal cells, contributing fundamentally to host innate immune efficacy^95^.

While *in vivo*, phosphate deprivation and oxidative attack on *C. albicans* cells occur together during their transition period from commensal to pathogenic lifestyle, we here sought to isolate the effects of these stressors on the cell wall as well as characterize their combined effect on the cell wall. We first defined in molecular detail the changes in cell wall architecture induced by phosphate deprivation, as the cell seeks to preserve precious phosphate for essential functions like nucleic acid biosynthesis and core carbon metabolism. We then exposed cells unable to import sufficient phosphate, the phosphate transport mutants, versus phosphate-replete wildtype cells to a sublethal H_2_O_2_ concentration, examining the effect of combined phosphate deprivation and peroxide stress.

Our findings extend current models by showing that ROS susceptibility is strongly shaped by cell-wall accessibility and architecture, not only by intracellular antioxidant capacity. We identify phosphate availability as a major determinant of wall structure and oxidative resilience, showing that phosphate homeostasis shapes not only intracellular metabolism but also the organization and dynamics of the extracellular wall. By demonstrating that the mobility, exposure, and redistribution of polysaccharides govern ROS interactions, we establish the cell wall as an active participant in oxidative protection and show that nutrient availability acts as a structural regulator of ROS resistance.

By analyzing intact *C. albicans* cells, we show that severe restriction of inorganic phosphate uptake extensively remodels the rigid wall core: the β-1,3-glucan-dominated scaffold characteristic of wildtype cells is replaced by a composite enriched in chitin, β-1,6-glucan, and mannan sidechains and exhibits reduced hydration and molecular mobility. This remodeling resembles stress-induced reinforcement observed under echinocandin or osmotic challenge, revealing that nutrient limitation alone can reconfigure the rigid scaffold and establishing phosphate availability as a structural regulator of the fungal cell wall. Phosphate-dependent changes in wall architecture also govern susceptibility to ROS. Hydrogen peroxide primarily targets the highly mobile, surface-exposed mannans and β-1,6-glucans, whereas the rigid core remains comparatively protected. In wildtype cells, oxidative exposure drives some mobile polymers into the rigid scaffold, reinforcing the wall; this response is markedly diminished in phosphate-transport mutants. Under the moderate concentration used in this study, peroxide-induced radicalization of mobile polysaccharides is found to be reversible, consistent with a transient redox-buffering mechanism, and H_2_O_2_ lacks intrinsic polymer specificity, reacting instead with solvent-exposed and dynamic sites. Together, these findings establish phosphate transport and cell-wall architecture as key determinants of oxidative susceptibility and identify accessibility, rather than monomer identity, as the principal regulator of ROS interaction in fungal cell walls.

High-resolution solid-state NMR thereby enables us to build an integrated structural model of both strains and to visualize how phosphate availability and ROS exposure reshape the cell-wall architecture. In the wildtype, the rigid core is organized primarily around β-1,3-glucan and a small portion of chitin, with additional contributions from a low content of β-1,6-glucan, and remains relatively hydrated and dynamic (**Fig. 5A**). In contrast, the rigid core of the phosphatetransport-deficient mutant is more heterogeneous, incorporating not only β-1,3-glucan and an increased amount of chitin, but also more α-1,2-mannan and β-1,6-glucan, which become associated with these rigid components (**Fig. 5B**). The reduced content of β-1,3-glucan and the increased abundance of chitin microfibrils lower hydration and restrict polymer mobility within the rigid fraction, resulting in a less accessible and more static architecture.

**Figure 5.**
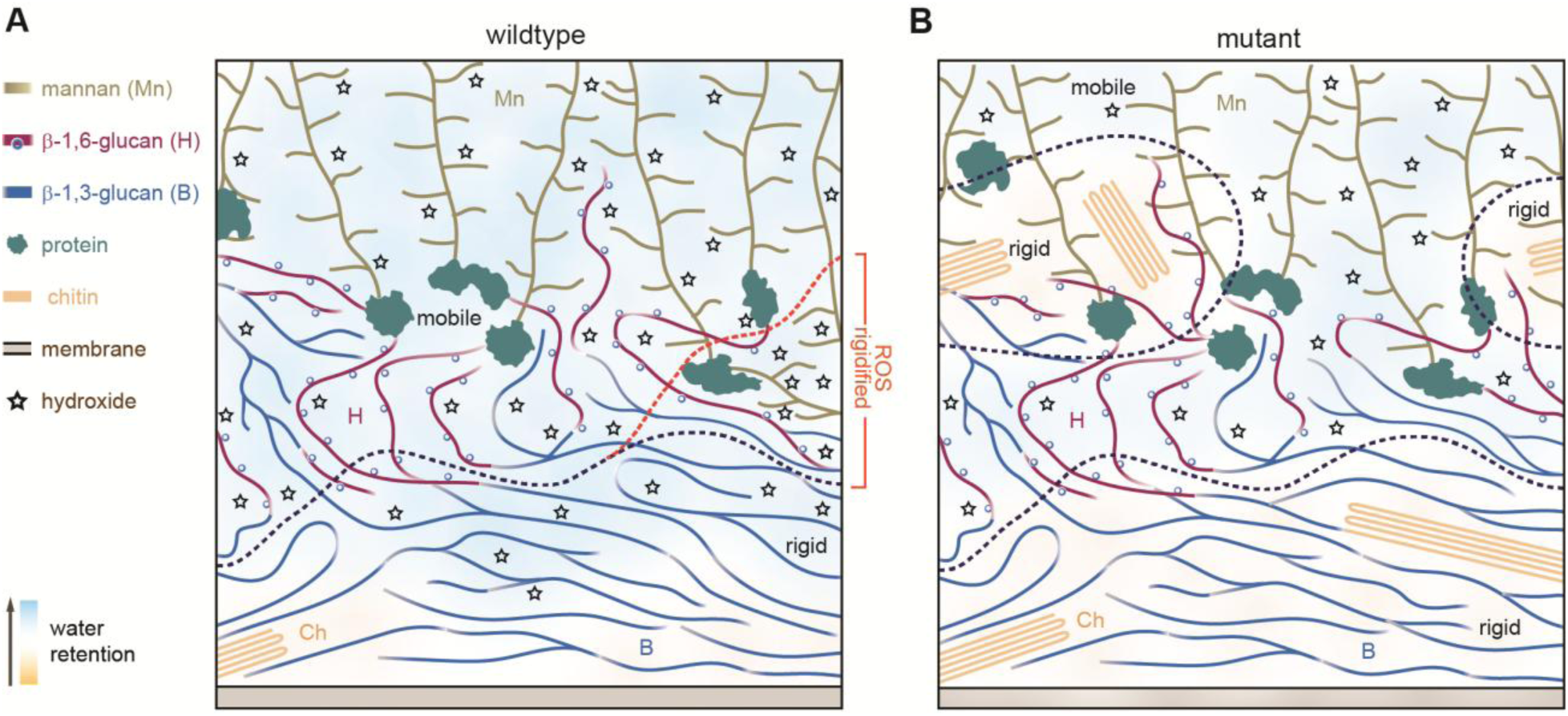
Schematics of wildtype and mutant C. albicans cell walls and peroxide retention. (**A**) Organization of the wildtype cell wall showing distinct inner and outer domains (separated by black dashed lines). Blue ribbon: β-1,3-glucan; purple ribbon: β-1,6-glucan; yellow branches: mannan; orange fibrils: chitin; gray: plasma membrane; green: protein; open blue circles: β-1,3-glucan branching from β-1,6-glucan backbone; black stars: retained peroxides. The orange-to-blue background gradient indicates increasing hydration. The orange dashed line marks the region that rigidifies upon hydroxide treatment, where hydroxides preferentially accumulate. (**B**) Schematic of a phosphate-transport-deficient mutant. Increased chitin content introduces additional rigid domains, causing normally flexible components (e.g., mannan sidechains and β-1,6-glucan) to become stiff, decreasing overall hydration and reducing the cell wall’s capacity to retain peroxides. Schematics reflect relative molecular composition but may not be strictly to scale.

Upon H_2_O_2_ exposure, mobile α-1,2-mannan and β-1,6-glucan in the wildtype are recruited into the rigid scaffold (**Fig. 5A**). Preferential broadening of α-1,2-mannan resonances in this newly rigidified domain indicates direct interaction between H_2_O_2_ and these polysaccharides. Oxidative damage involves both rigid and mobile fractions but occurring more extensively within the latter. In the mutant, H_2_O_2_ has limited access to the rigid core because of reduced hydration and increased packing, consistent with minimal broadening of rigid-core signals (**Fig. 5B**). Oxidative interactions occur primarily in the mobile layer, which rapidly loses NMR visibility in both strains, in agreement with molecular dynamics simulations showing that H_2_O_2_ is not intrinsically selective for any particular polymer. Together, these observations support a structural concept in which the wildtype wall intercepts ROS more effectively by engaging a substantial mobile carbohydrate fraction, a portion of the original rigid core, and polysaccharides that become rigid upon ROS exposure. In contrast, the mutant relies predominantly on its mobile carbohydrates, a significant fraction of which is already rigidified through interactions with β-1,3-glucan and chitin microfibrils and is therefore inaccessible to ROS. This limited capacity to redistribute and trap ROS contributes to the oxidative stress hypersensitivity of phosphate-transport mutants^13,25^.

The structural differences we observe are consistent with prior evidence that phosphate acquisition is closely linked to cell wall biosynthesis in *C. albicans*. Inorganic phosphate import supports intracellular phosphate pools and drives the production of nucleotide-sugar precursors required for β-glucan, chitin, and mannan synthesis^25,31^. Deletion of *PHO84* or disruption of the phosphate transport regulatory network increases susceptibility to cell-wall-active agents and compromises wall assembly, indicating that phosphate availability regulates wall architecture through its control of biosynthetic capacity^25,31^. Thus, the extensive remodeling we detect by solid-state NMR represents the structural outcome of phosphate-regulated biosynthetic pathways, extending beyond signaling and metabolic changes.

Our findings also reveal an unexpected biochemical intersection between phosphate metabolism and oxidative stress defense. Deletion of high-affinity phosphate transporters impairs intracellular phosphate homeostasis and attenuates TORC1 signaling, both of which are established regulators of oxidative adaptation and cell-wall integrity^13,25,31,53,96^. As a result, the phosphate transport mutant fails to initiate stress-induced redistribution of polysaccharides that is needed for reinforcing the rigid core of the wall. This indicates that phosphate sensing coordinates intracellular antioxidant responses with extracellular structural defenses. Reduced hydration and accessibility in the mutant further restrict peroxide binding and radical quenching. Thus, phosphate availability emerges as a central regulator that integrates nutrient sensing, redox homeostasis, and the mechanical architecture of the fungal cell wall.

Because Pho84-dependent phosphate acquisition and its downstream signaling pathways are conserved among fungal pathogens and required for virulence, targeting phosphate uptake or signaling could restrict the ability of pathogens to remodel and reinforce their cell wall during immune attack^53,97^. The pronounced vulnerability of the mannan-rich outer layer also identifies a structurally exposed domain that may be exploited pharmacologically or by host immune mechanisms. Together, our findings highlight phosphate homeostasis and wall architecture as actionable determinants of oxidative stress resilience and suggest new therapeutic approaches that combine nutrient limitation, cell wall-active agents, and fungal-specific oxidative stress^98^ to safely and effectively eradicate *C. albicans* infections even in immunocompromised hosts.

## METHODS

### Preparation of ^13^C-labeled yeast cells

The wildtype (JKC 915) and phosphate transport-deficient mutant (JKC 2830; *pho84Δ/Δ pho89Δ/Δ pho87Δ/Δ fgr2Δ/Δ*) of *C. albicans* were first pre-cultured in 50 mL of standard yeast nitrogen base (YNB) medium containing phosphate to ensure robust initial growth. The YNB medium was composed of 0.34 g (0.67%) YNB with ammonium sulfate and without amino acids, supplemented with 1 g of glucose (2% w/v) and 0.25 g of ammonium sulfate (5 g/L final concentration). A small inoculum (approximately the size of a pinhead) was transferred into 50 mL of this medium and incubated at 30°C with agitation at 200 rpm for approximately 15 h. Cells from this culture were subsequently diluted 1:20 by transferring 2.5 mL into fresh 50 mL YNB medium lacking phosphate but containing 2% glucose and ammonium sulfate, and incubated under the same shaking and temperature conditions for 24 h. This phosphate-deprivation step was repeated by pelleting cells via centrifugation at 1500 rpm for 15 min, decanting the supernatant, and resuspending the pellet in freshly autoclaved YNB medium without phosphate, containing 2% glucose and ammonium sulfate, followed by another 24 h incubation under identical conditions. For isotopic labeling, the cells were harvested again as above and transferred into 50 mL of freshly prepared YNB medium devoid of phosphate but supplemented with 2% uniformly labeled ^13^C glucose (1 g), 0.335 g of YNB without phosphate, and 3 mM of KH_2_PO_4_ (20.4 mg). The culture was incubated at 30°C with shaking at 200 rpm for 24 h to allow for incorporation of ^13^C into cellular biomass. Cells were subsequently harvested and packed into 3.2-mm MAS rotors for solid-state NMR analysis.

### Preparation of ^13^C yeast cells for evaluating oxidative stress response via solid-state NMR

Prior to detailed NMR analysis, pilot experiments were conducted to determine the concentrations of H_2_O_2_ that diminish growth in the strains used here, without fully inhibiting it. For solid-state NMR analysis, *C. albicans* cells were grown in 0.67% yeast nitrogen base (YNB with phosphate), 2% glucose, and 5 g/L ammonium sulfate in 100 mL milli-Q water. Cells from YPD plates were inoculated into fresh medium to a starting OD_600_ of approximately 0.5 and grown overnight at 30 °C with shaking at 200 rpm. Cultures then were incubated at the same conditions until the OD_600_ of the control reached 1.0 (approximately 15 h), at which point all cultures were harvested by centrifugation at 1500 rpm in falcon tubes. Pellets were washed with Milli-Q water to remove residual medium.

For oxidative stress experiments, cells were cultured in defined medium supplemented with uniformly labeled ^13^C-glucose as the primary carbon source. The inclusion of 0.8 mM H_2_O_2_ (30% stock solution) allowed for controlled induction of peroxide stress under defined nutrient conditions. Two experimental media containing 0.8 mM H_2_O_2_ and a control without H_2_O_2_ were prepared. Following overnight pre-growth, cells were inoculated into each condition to a standardized OD_600_ of 0.2 and incubated at 30 °C with shaking. Cell density in the control condition was monitored, and all cultures were harvested when the control reached an OD _600_ of 0.4. Pellets were washed with Milli-Q water to remove extracellular metabolites and media components. Natively hydrated cells, approximately 45-50 mg per sample, were packed into 3.2-mm MAS rotors (HZ16916, Cortecnet) for subsequent solid-state NMR analysis.

### 13C Solid-state NMR experiments

A variety of 1D and 2D ^13^C solid-state NMR experiments were conducted on two Bruker Avance Neo spectrometers at the MSU Max T. Rogers NMR Facility. High-resolution 1D and 2D spectra were acquired on the 800 MHz (18.8 T) spectrometer, while relaxation and water-editing experiments were performed on the 400 MHz (9.4 T) system. All spectra were acquired using 3.2-mm HCN probes at 15 kHz MAS and 293 K. ^13^C chemical shifts were externally referenced to the adamantane CH_2_ peak at 38.48 ppm on the tetramethylsilane (TMS) scale. Unless otherwise specified, typical radiofrequency field strengths were 80-100 kHz for ¹H decoupling, 62.5 kHz for ^1^H hard pulses, and 50-62.5 kHz for ^13^C. A summary of experimental parameters is provided in **Supplementary Table 1**. All spectra were processed using TopSpin 4.3.0, graphs were generated with OriginPro 9.0, and figures were prepared in Adobe Illustrator 2023 (v27.2.0).

1D ^13^C spectra were collected using different polarization schemes to selectively probe rigid and mobile components (**Supplementary Fig. 1**). Rigid components were detected by ^13^C cross-polarization (CP) with a 1 ms contact time^99,100^. Mobile components were measured using ^13^C direct polarization (DP) with short 2 s recycle delays, while extending the delay to 35 s enabled quantitative detection of all carbon species in the same DP experiment. Highly mobile and solvent-exposed molecules were identified using J-coupling-based refocused Insensitive Nuclei Enhancement by Polarization Transfer (INEPT) experiments^101^.

Resonance assignments for rigid molecules were obtained from 2D ^13^C-^13^C CORD homonuclear correlation spectra with a 53 ms mixing time, which detect predominantly intramolecular cross-peaks^102^. For mobile components, 2D DP refocused J-INADEQUATE spectra were measured to determine through-bond connectivity, with each of the four τ delays optimized to 2.3 ms^103^. Assigned chemical shifts were compared with entries in the Complex Carbohydrate Magnetic Resonance Database (CCMRD) to validate carbohydrate identities^104^. The ^13^C chemical shifts of carbohydrates identified in *C. albicans* are listed in **Supplementary Table 2**.

Water accessibility of polysaccharides was assessed using a water-edited 2D ^13^C-^13^C correlation experiment^105,106^. The experiment began with ¹H excitation followed by a ^1^H-T_2_ filter (1.4 ms × 2 for the wildtype sample and 1.5 ms × 2 for the phosphate transport mutant), which suppressed approximately 95% of carbohydrate signals while retaining 75-80% of the water magnetization (**Supplementary Fig. 3**). The remaining water magnetization was transferred to nearby polysaccharides through a 4 ms ^1^H-^1^H mixing period and subsequently to ¹³C via a 1 ms ^1^H-^13^C CP step for high-resolution ^13^C detection. A 50 ms DARR mixing period was used for both the water-edited spectrum and a control 2D spectrum with full signal intensity. The ratio of intensities between the water-edited spectrum (S) and the control spectrum (S_0_) was calculated, with S/S_0_ values reflecting the relative hydration levels at individual carbon sites. All intensities were pre-normalized to the number of scans acquired for each spectrum.

The dynamics of cell wall polysaccharides was examined using ^13^C spin-lattice (T_1_) relaxation, which was measured using a series of z-filter delays ranging from 0.1 μs to 12 s (**Supplementary Fig. 4a** and **Supplementary Table 3**) ^107^. For each well-resolved peak, the decrease in signal intensity with increasing z-filter time was quantified and fitted to a single-exponential equation to obtain the ^13^C-T_1_ relaxation constant for each carbon site. Peak intensities were pre-normalized by the number of scans. In addition, ^13^C-detected ^1^H-T_1ρ_ relaxation was measured using a Lee-Goldburg (LG) spin-lock sequence combined with LG-CP to suppress ^1^H spin diffusion during both the spin-lock and cross-polarization periods. This approach enabled site-specific analysis of ^1^H-T_1ρ_ relaxation through detection of the directly bonded ^13^C nuclei. The decay of peak intensity was fitted to a single-exponential function to obtain the ^1^H-T_1ρ_ relaxation time constant for each resolved site (**Supplementary Fig. 4b** and **Supplementary Table 3**).

The molecular composition was estimated by analyzing the intensities of distinct carbohydrate signals resolved in 2D ^13^C spectra. The composition of rigid components was estimated from 2D ^13^C-^13^C CORD spectra, and that of mobile molecules was evaluated using ^13^C DP refocused J-INADEQUATE spectra. Only well-resolved peaks were included in the analysis to mitigate uncertainty from spectral overlap. Relative polysaccharide abundances were determined by summing the integrated peak volumes for each polysaccharide and normalizing by the number of associated cross-peaks. The standard error for each polysaccharide was calculated as the standard deviation of the integrated peak volumes divided by the corresponding cross-peak count. The overall standard error was obtained as the square root of the sum of the squared individual standard errors. Percentage error was estimated by dividing each polysaccharide’s standard error by its mean integrated peak volume, and then scaling by its relative abundance and its contribution to the total integrated volume.

### 1H-detected solid-state NMR experiments

The mobile polysaccharide components of the *C. albicans* cell wall were analyzed in both the wildtype and phosphate transport-deficient strains using scalar-coupling-based ^1^H-detected solid-state NMR experiments. All measurements were performed on a Bruker Avance Neo 800 MHz spectrometer equipped with a 3.2-mm HCN triple-resonance MAS probe, operating at a spinning frequency of 15 kHz. The feasibility of this experiment at slow MAS arises from the exclusive detection of mobile molecules, whose rapid dynamics sufficiently attenuate ^1^H-^1^H dipolar couplings to permit high-resolution spectra. ^13^C chemical shifts were calibrated to the TMS scale using the methylene resonance of adamantane at 38.48 ppm, and ^1^H chemical shifts were referenced to DSS (trimethylsilylpropanesulfonate) in D_2_O, with the DSS peak set to 0 ppm.

Proton-detected 2D ^1^H-^13^C *J*-hCcH TOCSY (total correlation spectroscopy)^67^ and 3D ^1^H-^13^C *J*-hCCH TOCSY experiments were performed using DIPSI-3 (decoupling in the presence of scalar interactions) mixing^68^ to enable efficient magnetization transfer through scalar couplings. In this pulse-sequence notation, capital letters denote nuclei whose frequencies are evolved (thus generating corresponding ^13^C or ^1^H dimensions in the spectra), while lowercase letters indicate nuclei for which frequency evolution is omitted. The 90° pulse lengths were 3.5 µs (71.4 kHz) for ^1^H and 5 µs (50 kHz) for ^13^C. SPINAL-64 (small phase incremental alternation with 64 steps) heteronuclear decoupling^69^ was applied during both t_1_ and t_2_ periods with an radiofrequency field strength of 71.4 kHz, and WALTZ-16 (wideband alternating-phase low-power technique for zero-residual splitting) decoupling^70^ was used during ¹H acquisition with a field strength of 17 kHz. Water suppression was implemented using the MISSISSIPPI pulse sequence (multiple intense solvent suppression intended for sensitive spectroscopic investigation of protonated proteins)^71^, with a pulse duration of 40 ms and a radiofrequency field strength of 26 kHz. DIPSI-3 mixing for homonuclear ^13^C-^13^C TOCSY transfers employed a 2 ms spin-lock pulse followed by a 25.5 ms mixing period, both applied at a radiofrequency field strength of 17 kHz.

For the wildtype sample, a 3D *J*-hCCH TOCSY dataset was collected using 128 complex points in both the t_1_ and t_2_ dimensions, 8 scans per increment, and a 2 s recycle delay, resulting in a total acquisition time of 77 h 16 min. A corresponding 2D *J*-hCcH TOCSY spectrum was also recorded using identical parameters, except with one t_1_ point, 512 t_2_ points, and 32 scans per increment, yielding a total duration of 2 h 25 min. All spectra were acquired using States-TPPI for quadrature detection in the indirect dimensions^72^. The assigned ^13^C and ^1^H chemical shifts are provided in **Supplementary Table 4**. Complete acquisition parameters are summarized in **Supplementary Table 5**.

### Molecular Dynamics Simulations

Molecular dynamics simulations were performed to assess the propensity of hydrogen peroxide molecules to interact with specific glycans or their linkages, as illustrated in **Supplementary Fig. 11**. Four glycan chains were examined. A short chitin chain was constructed using ChitinBuilder^108^. A mannose chain and two mixed glucan chains with distinct linkage patterns were generated using psfgen in VMD (version 1.9.4a58)^109^, and subsequently minimized with LongBondEliminator^110^ to remove potential ring penetrations. Each glycan was solvated in a TIP3P water box with 6 nm side lengths^111^. To introduce oxidative conditions, 67 water molecules were replaced with hydrogen peroxide molecules, resulting in an effective hydrogen peroxide concentration of approximately 0.55 M. This high concentration allows for sufficient sampling in a limited simulation length.

Each system was simulated five times for 1000 ns using NAMD 3.0.1^112^ and the CHARMM36 force field for carbohydrates and chitin^113,114^. The simulations were performed at 298 K and 1 atm, maintained by a Langevin thermostat^115^ and isotropic barostat^116^. Each simulation step was 2 fs long, enabled by the SETTLE algorithm^117^. Long range electrostatics was handled by particle mesh Ewald^118^ summation with a 1.2 Å grid spacing. As is typical for CHARMM36, we use a 12 Å cutoff, smoothly switching at 10 Å.

The primary analysis for these trajectories was to analyze the interaction propensity for specific moieties on the glycans with hydrogen peroxide. We do this through evaluating a contact number that smoothly rather than harshly cuts off if atoms move further apart:

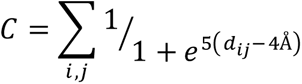

This contact definition has been used previously to quantify intermolecular interactions in plant biopolymers^119–121^. We evaluate this contact number over just heavy-atom pairs, which means that even a single contact is quite significant.

## DATA AVAILABILITY

All relevant data that support the findings of this study are provided in the article and Supplementary Information. All the original ssNMR data files will be deposited in the Zenodo repository, and the access code and DOI will be provided. Source Data are provided as a source data file.

## AUTHOR CONTRIBUTIONS

A.J., J.R.Y. and A.K. conducted solid-state NMR experiments and data analysis. W.Q. and A.J. prepared fungal samples. M. B. and J.V.V. performed MD simulations. A.J wrote the first draft of the manuscript. J.R.K and T.W. designed and supervised the project. All authors contributed to the manuscript writing.

## ACKNOWLEDGMENT

This study was primarily supported by National Institutes of Health (NIH) under award number R01AI173270 to T.W. The MD modeling was supported by the National Institute of General Medical Sciences of the National Institutes of Health under award number R35GM155317 to

J.V.V and supported through computational resources and services provided by the Institute for Cyber-Enabled Research at Michigan State University. *C. albicans* pilot experiments were supported by NIH NIAID under award number R21AI178693 to J.R.K.

## REFERENCES

1 Brown, G. et al. Hidden killers: human fungal infections. Sci. Transl. Med. 4, 165rv113 (2012).

2 Erwig, L. P. & Gow, N. A. Interactions of fungal pathogens with phagocytes. Nat. Rev. Microbiol. 14, 163–176 (2016).

3 Martinson, M. L. & Lapham, J. Prevalence of Immunosuppression Among US Adults. JAMA 331, 880–882 (2024).

4 Ghannoum, M. A. & Rice, L. B. Antifungal agents: mode of action, mechanisms of resistance, and correlation of these mechanisms with bacterial resistance. Clin. Microbiol. Rev. 12, 501–517 (1999).

5 Hoenigl, M. et al. The antifungal pipeline: fosmanogepix, ibrexafungerp, olorofim, opelconazole, and rezafungin. Drugs 81, 1703–1729 (2021).

6 Sun, S., Hoy, M. J. & Heitman, J. Fungal pathogens. Curr. Biol. 30, pR1163–R1169 (2020).

7 Pristov, K. E. & Ghannoum, M. A. Resistance of Candida to azoles and echinocandins worldwide. Clin. Microbiol. Infect. 25, 792–798 (2019).

8 Lamoth, F. Novel Therapeutic Approaches to Invasive Candidiasis: Considerations for the Clinician. Infect. Drug Resist. 16, 1087–1097 (2023).

9 Ksiezopolska, E. & Gabaldón, T. Evolutionary emergence of drug resistance in Candida opportunistic pathogens. Genes 9, 461 (2018).

10 Sanyaolu, A. et al. Candida auris: an overview of the emerging drug-resistant fungal infection. Infect. Chemother. 54, 236 (2022).

11 Arribas, V., Gil, C. & Molero, G. Deciphering the oxidative stressresponsein Candidaalbicans. Fungal Biol. Rev. 52, 100427 (2025).

12 Division, A. R. (ed World Health Organization) (World Health Organization, 2022).

13 Liu, N.-N. et al. Phosphate is the third nutrient monitored by TOR in Candida albicans and provides a target for fungal-specific indirect TOR inhibition. Proc. Natl. Acad. Sci. USA 114, 6346–6351 (2017).

14 Zara, V. et al. Yeast mitochondria lacking the phosphate carrier/p32 are blocked in phosphate transport but can import preproteins after regeneration of a membrane potential. Mol. Cell. Biol. 16, 6524–6531 (1996).

15 Bhalla, K. Q., X., Kretschmer, M. & Kronstad, J. W. The phosphate language of fungi. Trends Microbiol. 30, 338–349 (2022).

16 Pappas, P. G., Lionakis, M. S., Arendrup, M. C., Ostrosky -Zeichner, L. & Kullberg, B. J. Invasive candidiasis Nat. Rev. Dis. Primers 4, 18026 (2018).

17 Bays, D. J. & Savage, H. P. Candida albicans gastrointestinal colonization resistance: a host-microbiome balancing act. Infect. Immun. 93, e11610–11624 (2025).

18 White, S. J. et al. Self-regulation of Candida albicans population size during GI colonization. PLoS Pathog. 3, e184 (2007).

19 Tso, G. H. W. et al. Experimental evolution of a fungal pathogen into a gut symbiont Science 362, 589–595 (2018).

20. Koh, A. Y., Kohler, J. R., Coggshall, K. T., Rooijen, N. V. & Pier, G. B. Mucosal damage and neutropenia are required for Candida albicans dissemination PLOS Pathog. 4, e35 (2008).

21 Prochazkova, N. et al. Gut physiology and environment explain variations in human gut microbiome composition and metabolism. Nat. Microbiol. 9, 3210–3225 (2024).

22 Fine, K. D., Ogunji, F., Florio, R., Porter, J. & Ana, C. S. Investigation and diagnosis of diarrhea caused by sodium phosphate *Dig*. Dis. Sci. 43, 2708–2714 (1998).

23 King, W. R. et al. Glycerophosphocholine provision rescues Candida albicans growth and signaling phenotypes associated with phosphate limitation. Eukaryot. Cell 8, 00231–00223 (2023).

24 Rubio-Aliaga, I. & Krapf, R. Phosphate intake, hyperphosphatemia, and kidney function. *Eruo*. J. Physiol. 474, 935–947 (2022).

25 Acosta-Zaldívar, M. et al. Candida albicans’ inorganic phosphate transport and evolutionary adaptation to phosphate scarcity. PLoS Genet. 20, e1011156 (2024).

26 Lev, S. & Djordjevic, J. T. Why is a functional PHO pathway required by fungal pathogens to disseminate within a phosphate-rich host: A paradox explained by alkaline pH-simulated nutrient deprivation and expanded PHO pathway function. PLoS pathog. 14, e1007021 (2018).

27 Fradin, C. et al. Granulocytes govern the transcriptional response, morphology and proliferation of Candida albicans in human blood *Mol*. Microbiol. 56, 397–415 (2005).

28 Thewes, S. et al. In vivo and ex vivo comparative transcriptional profiling of invasive and non-invasive Candida albicans isolates identifies genes associated with tissue invasion. Mol. Microbiol. 63, 1606–1628 (2007).

29 Walker, L. A. et al. Genome-wide analysis of Candida albicans gene expression patterns during infection of the mammalian kidney. Fungal Genet. Biol. 46, 210–219 (2009).

30 Ikeh, M. et al. Pho4 mediates phosphateacquisition in Candidaalbicansand is vital for stressresistance and metal homeostasis. Mol. Biol. Cell 27, 2633–2801 (2016).

31 Liu, N.-N. et al. Phosphoric metabolites link phosphate import and polysaccharide biosynthesis for Candida albicans cell wall maintenance. mBio 11, 10.1128/mbio.03225-03219 (2020).

32 Molero, G. et al. The importance of the phagocytes’ innate response in resolution of the infection induced by a low virulent Candida albicans mutant. Scand. J. Immunol. 62, 224–233 (2005).

33 Destin, K. G., Linden, J. R., Laforce-Nesbitt, S. S. & Bliss, J. M. Oxidative burst and phagocytosis of neonatal neutrophils confronting Candida albicans and Candida parapsilosis. Early Hum. Dev. 85, 531–535 (2009).

34 da Silva Dantas, A., et al. Thioredoxin regulates multiple hydrogen peroxide-induced signaling pathways in Candida albicans. Mol. Cell. Biol. 30, 4550–4563 (2010).

35 Miramon, P. et al. A family of glutathione peroxidases contributes to oxidative stress resistance in Candida albicans. Med. Mycol. 52, 223–239 (2014).

36 Komalapriya, C. et al. Integrative model of oxidative stress adaptation in the fungal pathogen Candida albicans. PLoS One 10, e0137750 (2015).

37 Mavrianos, J., Desai, C. & Chauhan, N. Two-component histidine phosphotransfer protein Ypd1 is not essential for viability in Candida albicans. Eukaryot. Cell 13, 452–460 (2014).

38 Correia, I., Wilson, D., Hube, B. & Pla, J. Characterization of a Candida albicans mutant defective in all MAPKs highlights the major role of Hog1 in the MAPK signaling network. J. Fungi 6, 230 (2020).

39 Alonso-Monge, R. et al. The Hog1 mitogen-activated protein kinase is essential in the oxidative stress response and chlamydospore formation in Candida albicans. Eukaryot. Cell 2, 351–361 (2003).

40 Román, E., Correia, I., Prieto, D., Alonso, R. & Pla, J. The HOG MAPK pathway in Candida albicans: more than an osmosensing pathway. Int. Microbiol. 23, 23–29 (2020).

41 Eisman, B. et al. The Cek1 and Hog1 mitogen-activated protein kinases play complementary roles in cell wall biogenesis and chlamydospore formation in the fungal pathogen Candida albicans. Eukaryot. Cell 5, 347–358 (2006).

42 Gulshan, K., Thommandru, B. & Moye-Rowley, W. S. Proteolytic degradation of the Yap1 transcription factor is regulated by subcellular localization and the E3 ubiquitin ligase Not4. J. Biol. Chem. 287, 26796–26805 (2012).

43 Patterson, M. J. et al. Ybp1 and Gpx3 signaling in Candidaalbicansgovern hydrogen peroxide -induced oxidation of the Cap1 transcription factor and macrophage escape. Antioxid. Redox Signal. 19, 2244–2260 (2013).

44 Wang, Y. et al. Cap1p is involved in multiple pathways of oxidative stress response in Candida albicans. Free Radic. Biol. Med. 40, 1201–1209 (2006).

45 Gow, N. A., Latge, J.-P. & Munro, C. A. The fungal cell wall: structure, biosynthesis, and function. Microbiol. Spectr. 5, 10.1128/microbiolspec.funk-0035-2016 (2017).

46 Brown, G. D. et al. The pathobiology of human fungal infections. Nat. Rev. Microbiol. 22, 687–704 (2024).

47 Katsipoulaki, M. et al. Candida albicans and Candida glabrata: global priority pathogens. Microbiol. Mol. Biol. Rev. 88, e00021–00023 (2024).

48 Odds, F. C., Brown, A. J. & Gow, N. A. Antifungal agents: mechanisms of action. Trends Microbiol. 11, 272–279 (2003).

49 Hall, R. A. et al. The Mnn2 mannosyltransferase family modulates mannoprotein fibril length, immune recognition and virulence of Candida albicans. PLoS pathog. 9, e1003276 (2013).

50 Gow, N. A. & Lenardon, M. D. Architecture of the dynamic fungal cell wall. Nat. Rev. Microbiol. 21, 248–259 (2023).

51 Bekirian, C. et al. β-1, 6-glucan plays a central role in the structure and remodeling of the bilaminate fungal cell wall. Elife 13, RP100569 (2024).

52 Gow, N. A. R. Fungal cell wall biogenesis: structural complexity, regulation and inhibition. Fungal Genet. Biol. 179, 103991 (2025).

53 Liu, N.-N. et al. Intersection of phosphate transport, oxidative stress and TOR signalling in Candida albicans virulence. PLoS Pathog. 14, e1007076 (2018).

54 Yin, Z. et al. A proteomic analysis of the salt, cadmium and peroxide stress responses in Candida albicans and the role of the Hog1 stress-activated MAPK in regulating the stress-induced proteome. Proteomics 9, 4686–4703 (2009).

55 Enjalbert, B., MacCallum, D. M., Odds, F. C. & Brown, A. J. Niche-specific activation of the oxidative stress response by the pathogenic fungus Candida albicans. Infect. Immun. 75, 2143–2151 (2007).

56. Amador-García, A., et al. Extending the proteomic characterization of Candida albicans exposed to stress and apoptotic inducers through data-independent acquisition mass spectrometry. mSystems 6, e0094621 (2021).

57 Legrand, M., Chan, C. L., Jauert, P. A. & Kirkpatrick, D. T. Analysis of base excision and nucleotide excision repair in Candida albicans. Microbiology 154, 2446–2456 (2008).

58 Gautam, I. et al. Breaking down the wall: Solid-state NMR illuminates how fungi build and remodel diverse cell walls. PLoS Pathog. 21, e1013678 (2025).

59 Latge, J. P. & Wang, T. Modern biophysics redefines our understanding of fungal cell wall structure, complexity and dynamics. mBio 13, e01145–01122 (2022).

60 Reif, B., Ashbrook, S. E., Emsley, L. & Hong, M. Solid-State NMR Spectroscopy. Nat. Rev. Methods Primers 1, 2 (2021).

61 Dickwella Widanage, M. C., et al. Adaptative survival of Aspergillus fumigatus to echinocandins arises from cell wall remodeling beyond β− 1, 3-glucan synthesis inhibition. Nat. Commun. 15, 6382 (2024).

62 Chakraborty, A. et al. A molecular vision of fungal cell wall organization by functional genomics and solid-state NMR. Nat. Commun. 12, 6346 (2021).

63 Lamon, G. et al. Solid-state NMR molecular snapshots of Aspergillus fumigatus cell wall architecture during a conidial morphotype transition. Proc. Natl. Acad. Sci. USA 120, e2212003120 (2023).

64 Fernando, L. D. et al. Structural adaptation of fungal cell wall in hypersaline environment. Nat. Commun. 14, 7082 (2023).

65 Yarava, J. R., Gautam, I., Jacob, A., Fu, R. & Wang, T. Proton -Detected Solid-State NMR for Deciphering Structural Polymorphism and Dynamic Heterogeneity of Cellular Carbohydrates in Pathogenic Fungi. J. Am. Chem. Soc. 147, 17416–17432 (2025).

66 Bishoyi, A. K. et al. Solid-State NMR Reveals Reorganization of the Aspergillus fumigatus Cell Wall Due to a Host-Defence Peptide. Angew. Chem. Int. Ed. 64, e20259012 (2025).

67 Gautam, I. et al. Comparative analysis of polysaccharide and cell wall structure in Aspergillus nidulans and Aspergillus fumigatus by solid-state NMR. Carbohydr. Polym. 348, 122907 (2025).

68 Ehren, H. L. et al. Characterization of the cell wall of a mushroom forming fungus at atomic resolution using solid-state NMR spectroscopy. Cell Surf. 6, 100046 (2020).

69 Safeer, A. et al. Probing Cell-Surface Interactions in Fungal Cell Walls by High-Resolution 1H- Detected Solid-State NMR Spectroscopy. Chem. Eur. J. 29, e202202616 (2023).

70 Cheng, Q. et al. Molecular architecture of chitin and chitosan-dominated cell walls in zygomycetous fungal pathogens by solid-state NMR. Nat. Commun. 15, 8295 (2024).

71 Delcourte, L. et al. Magic-angle spinning NMR spectral editing of polysaccharides in whole cells using the DREAM scheme. Methods 230, 59–67 (2024).

72 Chatterjee, S., Prados-Rosales, R., Itin, B., Casadevall, A. & Stark, R. E. Solid-state NMR reveals the carbon-based molecular architecture of Cryptococcus neoformans fungal eumelanins in the cell wall. J. Biol. Chem. 290, 13779–13790 (2015).

73 Chrissian, C. et al. Solid-state NMR spectroscopy identifies three classes of lipids in Cryptococcus neoformans melanized cell walls and whole fungal cells. J. Biol. Chem. 295, 15083–15096 (2020).

74 Ankur, A. et al. Polymorphic α-Glucans as Structural Scaffolds in Cryptococcus Cell Walls for Chitin, Capsule, and Melanin: Insights From 13C and 1H Solid-State NMR. Angew. Chem. Int. Ed. 64, e202510409 (2025).

75 Lends, A. et al. Molecular Distinction of Cell Wall and Capsular Polysaccharides in Encapsulated Pathogens by In Situ Magic-Angle Spinning NMR Techniques. J. Am. Chem. Soc. 147, 6813–6824 (2025).

76 Jacob, A. et al. α-glucan remodeling by GH13-domain enzymes shapes fungal cell wall architecture. Proc. Natl. Acad. Sci. USA 122, e2505509122 (2025).

77 Dickwella Widanage, M. C., et al. Distinct echinocandin responses of Candida albicans and Candida auris cell walls revealed by solid-state NMR. Nat. Commun. 16, 6295 (2025).

78 Bahri, S. et al. 1H-detected characterization of carbon-carbon networks in highly flexible protonated biomolecules using MAS NMR *J*. Biomol. NMR 77, 111–119 (2023).

79 Nasution, O. et al. Hydrogen peroxide induces hyphal differentiation in Candida albicans. Eukaryot. Cell 7, 2008–2011 (2008).

80 Jamieson, D. J., Stephen, D. W. & Terrière, E. C. Analysis of the adaptive oxidative stress response of Candida albicans. FEMS Microbiol. Lett. 138, 83–88 (1996).

81 Fradin, C. et al. Granulocytes govern the transcriptional response, morphology and proliferation of Candida albicans in human blood. Mol. Microbiol. 56, 397–415 (2005).

82 Lorenz, M. C., Bender, J. A. & Fink, G. R. Transcriptional response of Candida albicans upon internalization by macrophages. Eukaryot. Cell 3, 1076–1087 (2004).

83 Bertini, I., Luchinat, C., Parigi, G. & Pierattelli, R. NMR spectroscopy of paramagnetic metalloproteins. ChemBioChem 6, 1536–1549 (2005).

84 Sengupta, I., Nadaud, P. S., Helmus, J. J., Schwieters, C. D. & Jaroniec, C. P. Protein fold determined by paramagnetic magic-angle spinning solid-state NMR spectroscopy. Nat. Chem. 4, 410–417 (2012).

85 Chen, X. et al. Free radical-mediated degradation of polysaccharides: Mechanism of free radical formation and degradation, influence factors and product properties. Food Chem. 365, 130524 (2021).

86 Schweikert, C., Liszkay, A. & Schopfer, P. Scission of polysaccharides by peroxidase-generated hydroxyl radicals. Phytochemistry 53, 565–570 (2000).

87 Hangasky, J. A., Iavarone, A. T. & Marletta, M. A. Reactivity of O2 versus H2O2 with polysaccharide monooxygenases. Proc. Natl. Acad. Sci. USA 115, 4915–4920 (2018).

88 Duan, J. & Kasper, D. L. Oxidative depolymerization of polysaccharides by reactive oxygen/nitrogen species. Glycobiology 21, 401–409 (2011).

89 Ofoedu, C. E. et al. Hydrogen peroxide effects on natural-sourced polysacchrides: free radical formation/production, degradation process, and reaction mechanism—a critical synopsis. Foods 10, 699 (2021).

90 Frohner, I. E., Bourgeois, C., Yatsyk, K., Majer, O. & Kuchler, K. Candida albicans cell surface superoxide dismutases degrade host-derived reactive oxygen species to escape innate immune surveillance. Mol. Microbiol. 71, 240–252 (2009).

91 Gleason, J. E. et al. Candida albicans SOD5 represents the prototype of an unprecedented class of Cu-only superoxide dismutases required for pathogen defense. Proc. Natl. Acad. Sci. USA 11, 5866–5871 (2014).

92 Bienert, G. P. & Chaumont, F. Aquaporin-facilitated transmembrane diffusion of hydrogen peroxide. Biochim. Biophys. Acta 1840, 1596–1604 (2014).

93 Lee, S., Chen, H., Yeh, Y., Xue, Y. & Lan, C. The Transcription Factor Sfp1 Regulates the Oxidative Stress Response in Candida albicans. Microorganisms 7, 131 (2019).

94 da Silva Dantas, A., et al. Oxidative Stress Responses in the Human Fungal Pathogen, Candida albicans. Biomolecules 25, 142–165 (2015).

95 Kos, I. et al. Mechanisms Underlying the Delayed Activation of the Cap1 Transcription Factor in Candida albicans following Combinatorial Oxidative and Cationic Stress Important for Phagocytic Potency. mBio 7, e00331–00316 (2016).

96 Köhler, J. R., Acosta-Zaldívar, M. & Qi, W. Phosphate in Virulence of Candida albicans and Candida glabrata. J. Fungi 6, 40 (2020).

97 Ikeh, M., Ahmed, Y. & Quinn, J. Phosphate Acquisition and Virulence in Human Fungal Pathogens Microorganisms 5, 48 (2017).

98 Qi, W. et al. Metabolic pathway competition sensitizes thioredoxin reductase-depleted Candida albicans to cell wall stress and antifungals. BioRxiv, DOI: 10.64898/2025.12.02.691982 (2025).

99 Pines, A., Gibby, M. G. & Waugh, J. S. Proton-enhanced nuclear induction spectroscopy 13C chemical shielding anisotropy in some organic solids. Chem. Phys. Lett. 15, 373–376 (1972).

100 Levitt, M. H., Suter, D. & Ernst, R. R. Spin dynamics and thermodynamics in solid-state NMR cross polarization. J. Chem. Phys. 84, 4243–4255 (1986).

101 Elena, B., Lesage, A., Steuernagel, S., Böckmann, A. & Emsley, L. Proton to carbon-13 INEPT in solid-state NMR spectroscopy. J. Am. Chem. Soc. 127, 17296–17302 (2005).

102 Hou, G., Yan, S., Trébosc, J., Amoureux, J. P. & Polenova, T. Broadband homonuclear correlation spectroscopy driven by combined R2(n)(v) sequences under fast magic angle spinning for NMR structural analysis of organic and biological solids. J. Magn. Reson. 232, 18–30 (2013).

103 Cadars, S. et al. The refocused INADEQUATE MAS NMR experiment in multiple spin-systems: interpreting observed correlation peaks and optimising lineshapes. J. Magn. Reson. 188, 24–34 (2007).

104 Kang, X. et al. CCMRD: a solid-state NMR database for complex carbohydrates. J. Biomol. NMR 74, 239–245 (2020).

105 White, P. B., Wang, T., Park, Y. B., Cosgrove, D. J. & Hong, M. Water-polysaccharide interactions in the primary cell wall of Arabidopsis thaliana from polarization transfer solid-state NMR. J. Am. Chem. Soc. 136, 10399–10409 (2014).

106 Ader, C. et al. Structural rearrangements of membrane proteins probed by water-edited solid-state NMR spectroscopy. J. Am. Chem. Soc. 131, 170–176 (2009).

107 Torchia, D. A. Measurement of proton-enhanced carbon-13 T1 values by a method which suppresses artifacts. J. Magn. Reson. 30, 613–616 (1978).

108 Malaspina, D. & Faraudo, J. Chitin Builder: a VMD tool for the generation of structures of chitin molecular crystals for atomistic simulations. J. Open Source Softw. 9, 5771 (2024).

109 Humphrey, W., Dalke, A. & Schulten, K. VMD: visual molecular dynamics. J. Mol. Graph. 14, 33–38 (1996).

110 Sarkar, D., Kulke, M. & Vermaas, J. V. LongBondEliminator: A molecular simulation tool to remove ring penetrations in biomolecular simulation systems. Biomolecules 13, 107 (2023).

111 Jorgensen, W. L., Chandrasekhar, J., Madura, J. D., Impey, R. W. & Klein, M. L. Comparison of simple potential functions for simulating liquid water. J. Chem. Phys. 79, 926–935 (1983).

112 Phillips, J. C. et al. Scalable molecular dynamics on CPU and GPU architectures with NAMD. J. Chem. Phys. 153 (2020).

113 Guvench, O., Hatcher, E., Venable, R. M., Pastor, R. W. & MacKerell Jr, A. D. CHARMM additive all-atom force field for glycosidic linkages between hexopyranoses. J. Chem. Theory Comput. 5, 2353–2370 (2009).

114 Guvench, O. et al. Additive empirical force field for hexopyranose monosaccharides. J. Comp. Chem. 29, 2543–2564 (2008).

115 Kubo, R. The fluctuation-dissipation theorem. Rep. Prog. Phys. 29, 255 (1966).

116 Feller, S. E., Zhang, Y., Pastor, R. W. & Brooks, B. R. Constant pressure molecular dynamics simulation: The Langevin piston method. J. Chem. Phys. 103, 4613–4621 (1995).

117 Miyamoto, S. & Kollman, P. A. Settle: An analytical version of the SHAKE and RATTLE algorithm for rigid water models. J. Comput. Chem. 13, 952–962 (1992).

118 Essmann, U. et al. A smooth particle mesh Ewald method. J. Chem. Phys. 103, 8577–8593 (1995).

119 Vermaas, J. V. et al. Mechanism of lignin inhibition of enzymatic biomass deconstruction. Biotechnol. Biofuels Bioprod. 8, 217 (2015).

120 Vermaas, J. V., Crowley, M. F. & Beckham, G. T. A quantitative molecular atlas for interactions between lignin and cellulose. ACS Sustain. Chem. Eng. 7, 19570–19583 (2019).

121 Sarkar, D. et al. Diffusion in intact secondary cell wall models of plants at different equilibrium moisture content. Cell Surf. 9, 100105 (2023).

