## Supplementary Material for "Oxidative Stress-Responsive Cell Wall Remodeling Depends on Phosphate in *Candida albicans*"

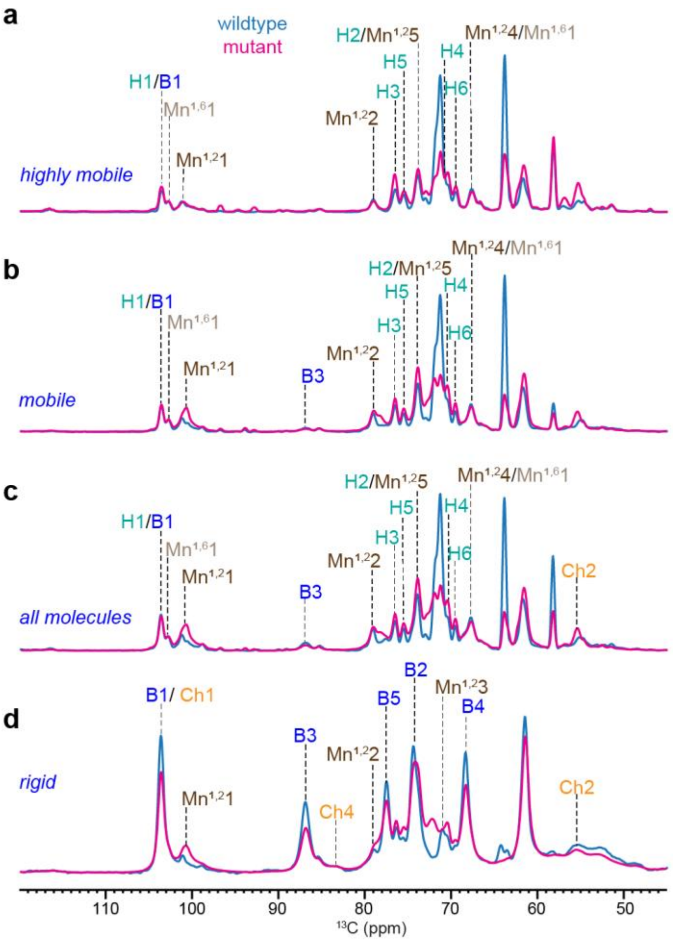


**Supplementary Figure 1. Dynamical gradient of polysaccharides in *Candida albicans* cell walls.** From top to bottom are four sets of 1D $^{13}$C spectra measured with (**a**) refocused INEPT experiment for probing the most dynamic molecules. (**b**) DP spectra with short recycle delay of 2 s for selection of mobile components. (**c**) DP with long recycle delays (35 s) for quantitative detection of all molecules. (**d**) CP for selecting rigid polysaccharides. The spectra of *C. albicans* wildtype and phosphate transport mutant are shown in blue and magenta, respectively. For example, the Mn$^{1,2}$1 peak at 101 ppm annotates the carbon 1 of a-1,2-mannose (Mn$^{1,2}$1).

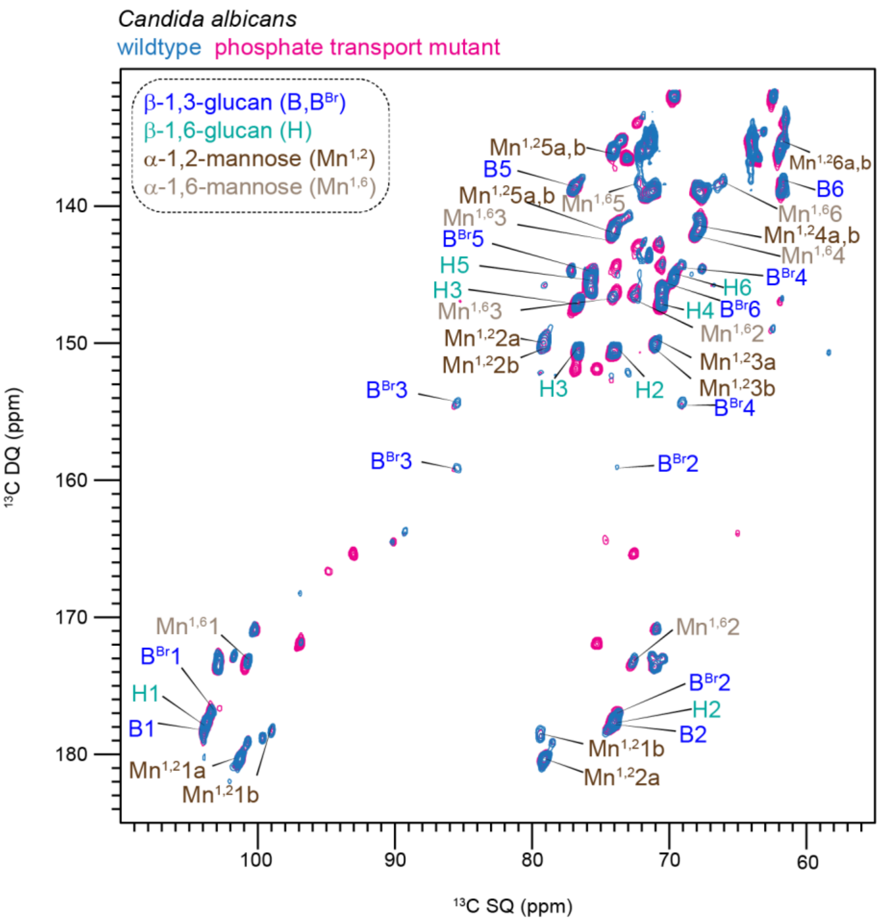


**Supplementary Figure 2. 2D $^{13}$C DP refocused J-INADEQUATE spectrum of *C. albicans* samples.** Signals are detected for linear β-1,3-glucan (B), linear β-1,6-glucan (H), β-1,3,6-glucopyranose unit ($B^{Br}$) of the branched glucan, as well as α-1,6-mannan ($Mn^{1,6}$) and α-1,2-mannose ($Mn^{1,2}$) residues of mannan. The changes noted are mainly from soluble, small molecules rather than these cell wall polymers.

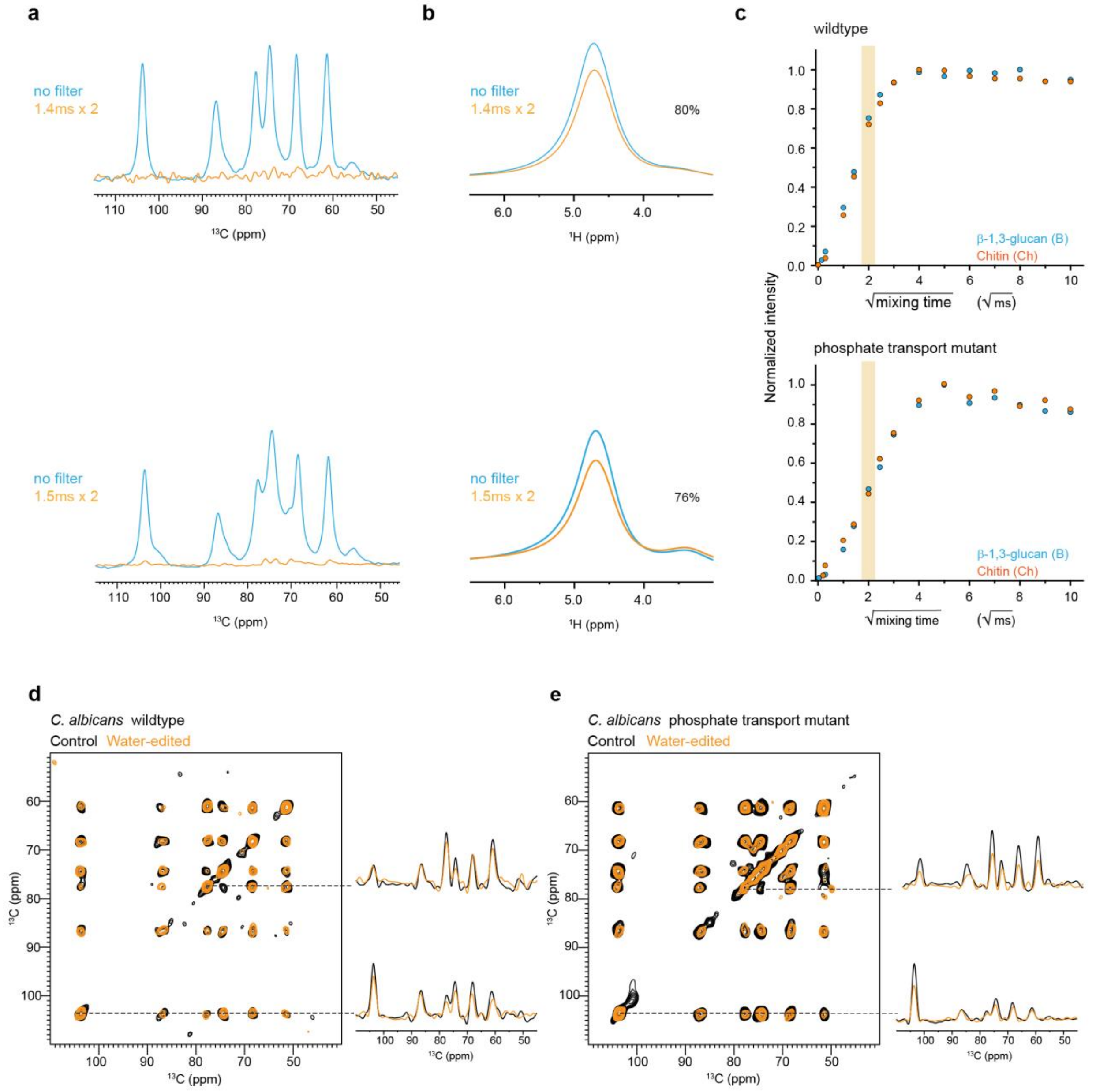


**Supplementary Figure 3. Water-edited spectra of *C. albicans* to access polymer hydration.** (**a**) $^{1}$H-T$_2$ filtered (orange) and control (cyan) $^{13}$C spectra are shown for wildtype (top) and quadruple mutant (bottom) *C. albicans* samples. No spin diffusion was applied. Approximately 95% of carbohydrate $^{13}$C signals were removed by the $^{1}$H-T$_2$ filter. (**b**) $^{1}$H-T$_2$ filtered (orange) and control (cyan) $^{1}$H NMR spectra, with 80% and 76% of water signal retained for wildtype and quadruple mutant respectively after the $^{1}$H-T$_2$ filter. (**c**) Representative water-to-polysaccharide $^{1}$H spin diffusion buildup curves are shown for wildtype (top) and quadruple mutant (bottom). Overlay of 2D water-edited (orange) and control (black) $^{13}$C-$^{13}$C correlation spectra of (**d**) wildtype sample and (**e**) quadruple mutant. Representative 1D slices extracted from the 2D $^{13}$C-$^{13}$C correlation spectra are shown for each sample. All spectra were measured on a 400 MHz spectrometer at 10 kHz MAS.

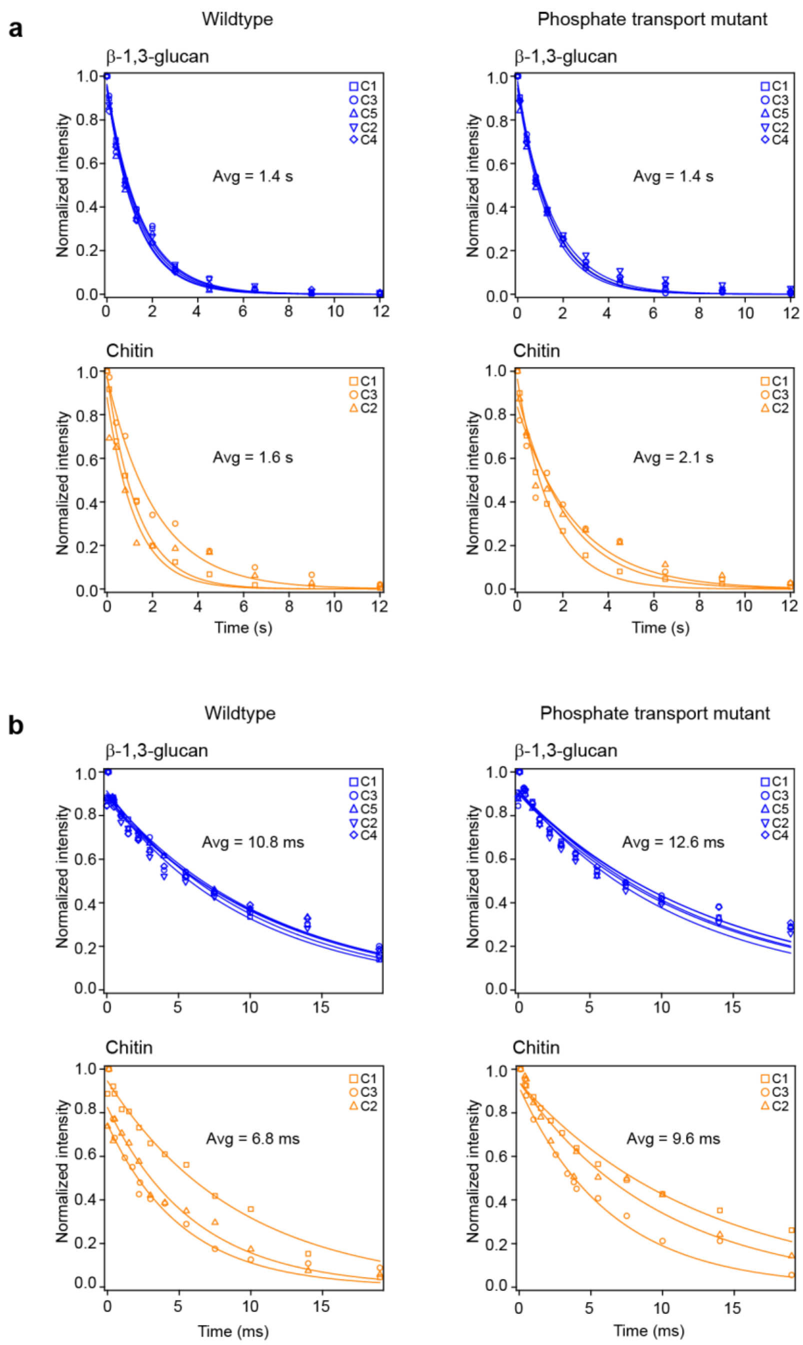


**Supplementary Figure 4**. **Relaxation curves of polysaccharides in *Candida* cell walls.** The relaxation decay curves are shown separately for (**a**) $^{13}$C T1 and (**b**) $^{1}$H T1ρ of the *C. albicans* wildtype and quadruple mutant.

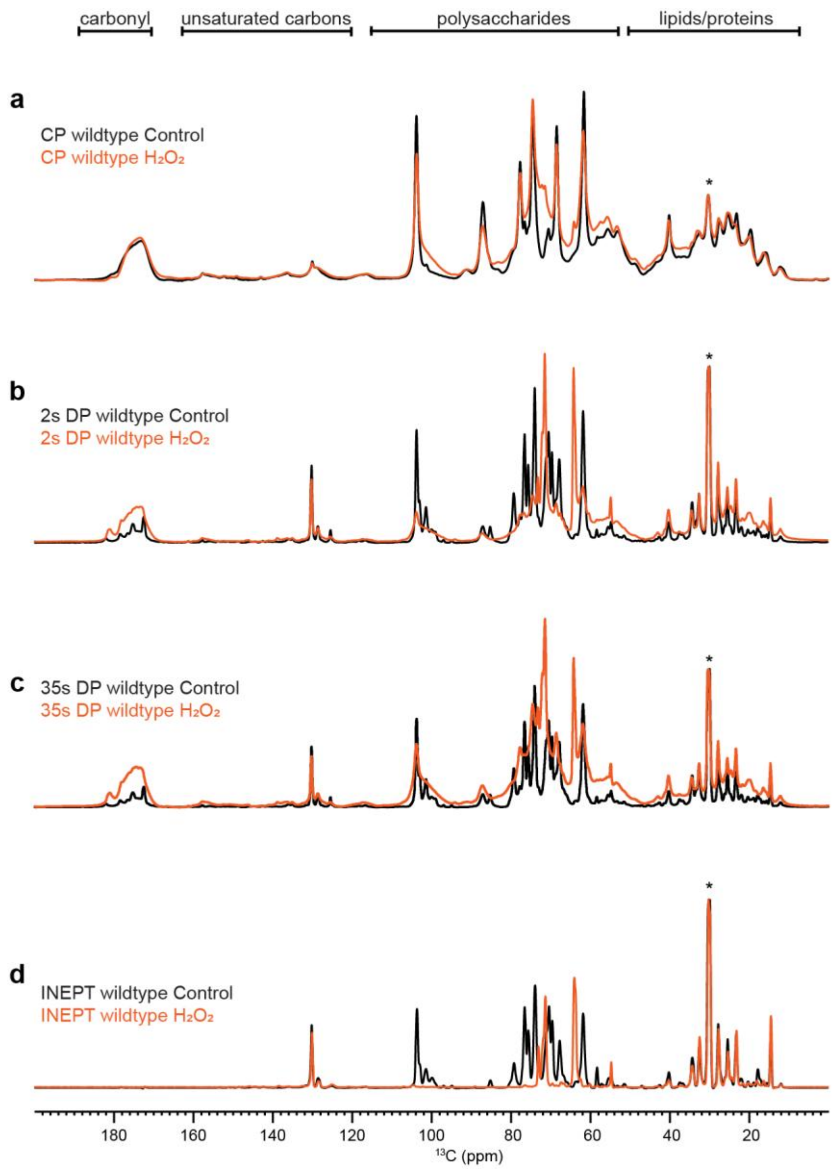


**Supplementary Figure 5. Full 1D $^{13}C$ spectra of wildtype *C. albicans* cell wall after $H_2O_2$ treatment.** From top to bottom are four sets of 1D $^{13}C$ spectra measured with (**a**) CP for selecting rigid polysaccharides. (**b**) DP spectra with a short recycle of 2s for selection of mobile components. (**c**) DP with long recycle delays (35s) for quantitative detection of all molecules. (**d**) Refocused INEPT experiments for probing the most dynamic molecules. The spectra for the wildtype of *C. albicans* control and $H_2O_2$ treated is shown in black and orange color, respectively.

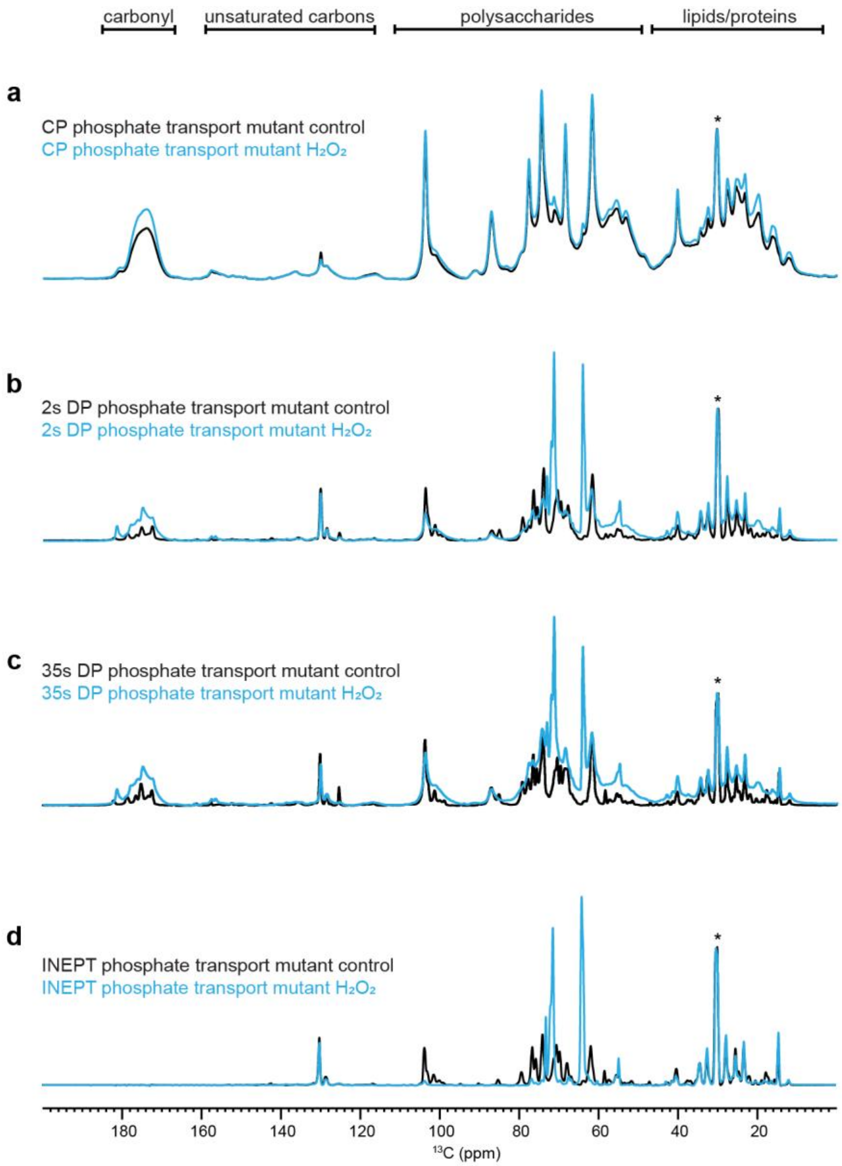


**Supplementary Figure 6. Full 1D $^{13}C$ spectra of mutant *C. albicans* cell wall after $H_2O_2$ treatment.** From top to bottom are four sets of 1D $^{13}C$ spectra measured with (**a**) CP for selecting rigid polysaccharides. (**b**) DP spectra with a short recycle of 2s for selection of mobile components. (**c**) DP with long recycle delays (35s) for quantitative detection of all molecules. (**d**) Refocused INEPT experiments for probing the most dynamic molecules. The spectra for the quadruple mutant of *C. albicans* control and $H_2O_2$ treated is shown in black and cyan color, respectively.

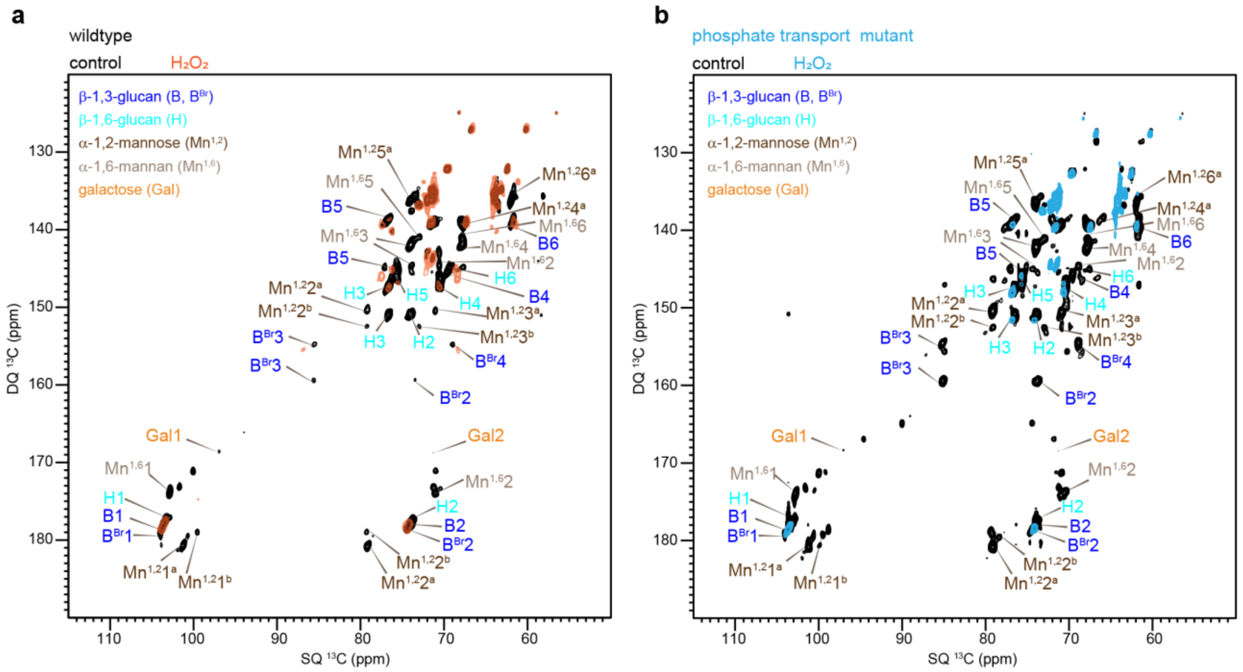


**Supplementary Figure 7**. **Full carbohydrate region of 2D $^{13}$C DP refocused J-INADEQUATE spectra.** Mobile molecules are detected in (**a**) wild-type and (**b**) mutant cells under control or $H_2O_2$-treated conditions.

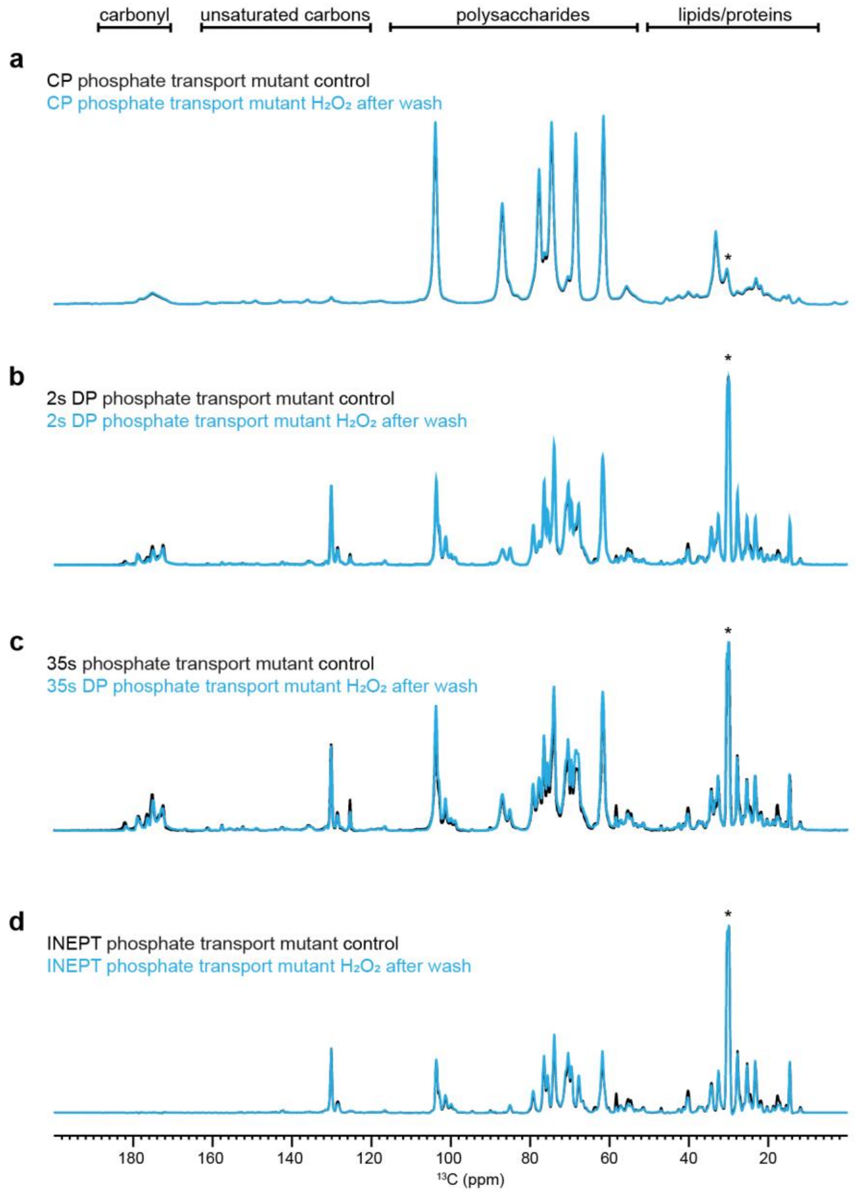


**Supplementary Figure 8. 1D $^{13}$C spectra of $H_2O_2$-treated mutant *C. albicans* cell wall after washing**. From top to bottom are four sets of 1D $^{13}$C spectra measured with (**a**) CP for selecting rigid polysaccharides. (**b**) DP spectra with a short recycle of 2s for selection of mobile components. (**c**) DP with long recycle delays (35s) for quantitative detection of all molecules. (**d**) Refocused INEPT experiments for probing the most dynamic molecules. The spectra for the quadruple mutant of *C. albicans* control and washed after $H_2O_2$ treatment is shown in black and cyan color, respectively.

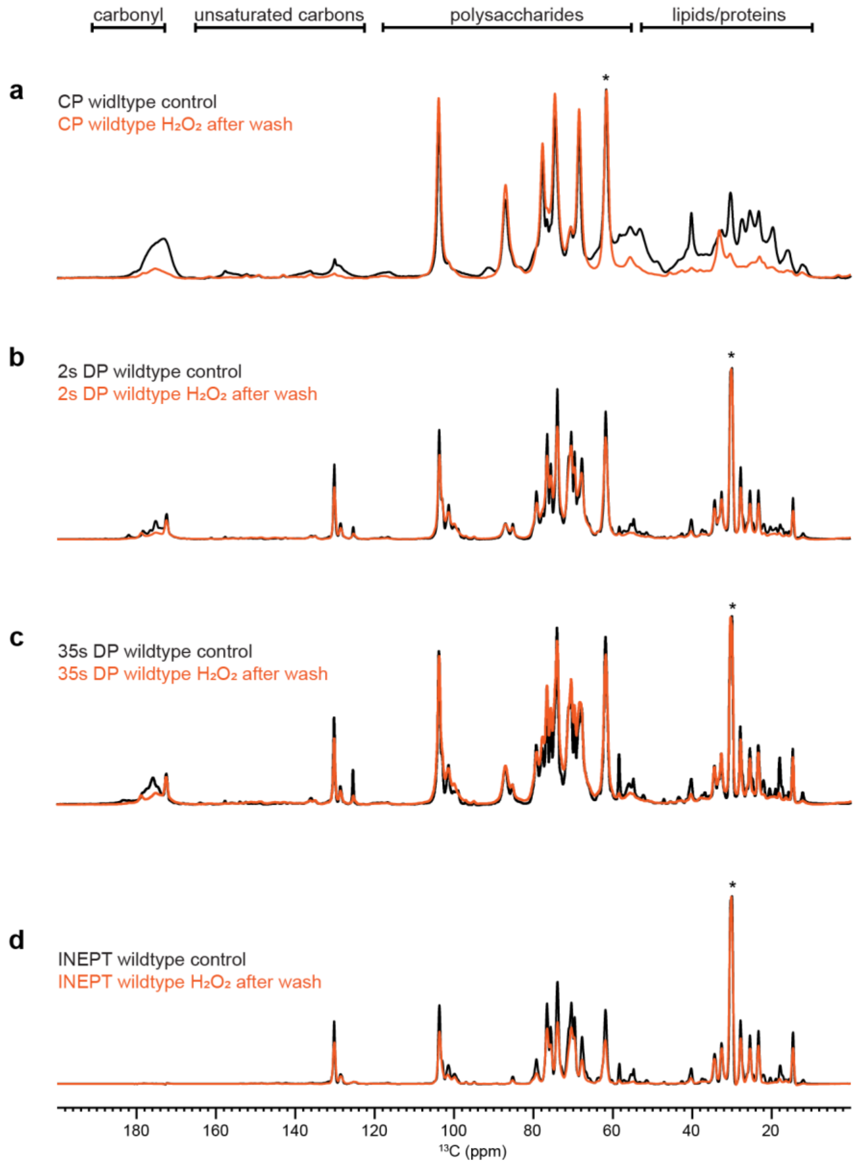


**Supplementary Figure 9. 1D $^{13}C$ spectra of $H_2O_2$-treated wildtype cell wall after washing.** From top to bottom are four sets of 1D $^{13}C$ spectra measured with (**a**) CP for selecting rigid polysaccharides. (**b**) DP spectra with a short recycle of 2s for selection of mobile components. (**c**) DP with long recycle delays (35s) for quantitative detection of all molecules. (**d**) Refocused INEPT experiments for probing the most dynamic molecules. The spectra for the wildtype of *C. albicans* control and washed after $H_2O_2$ treatment is shown in black and orange color, respectively.

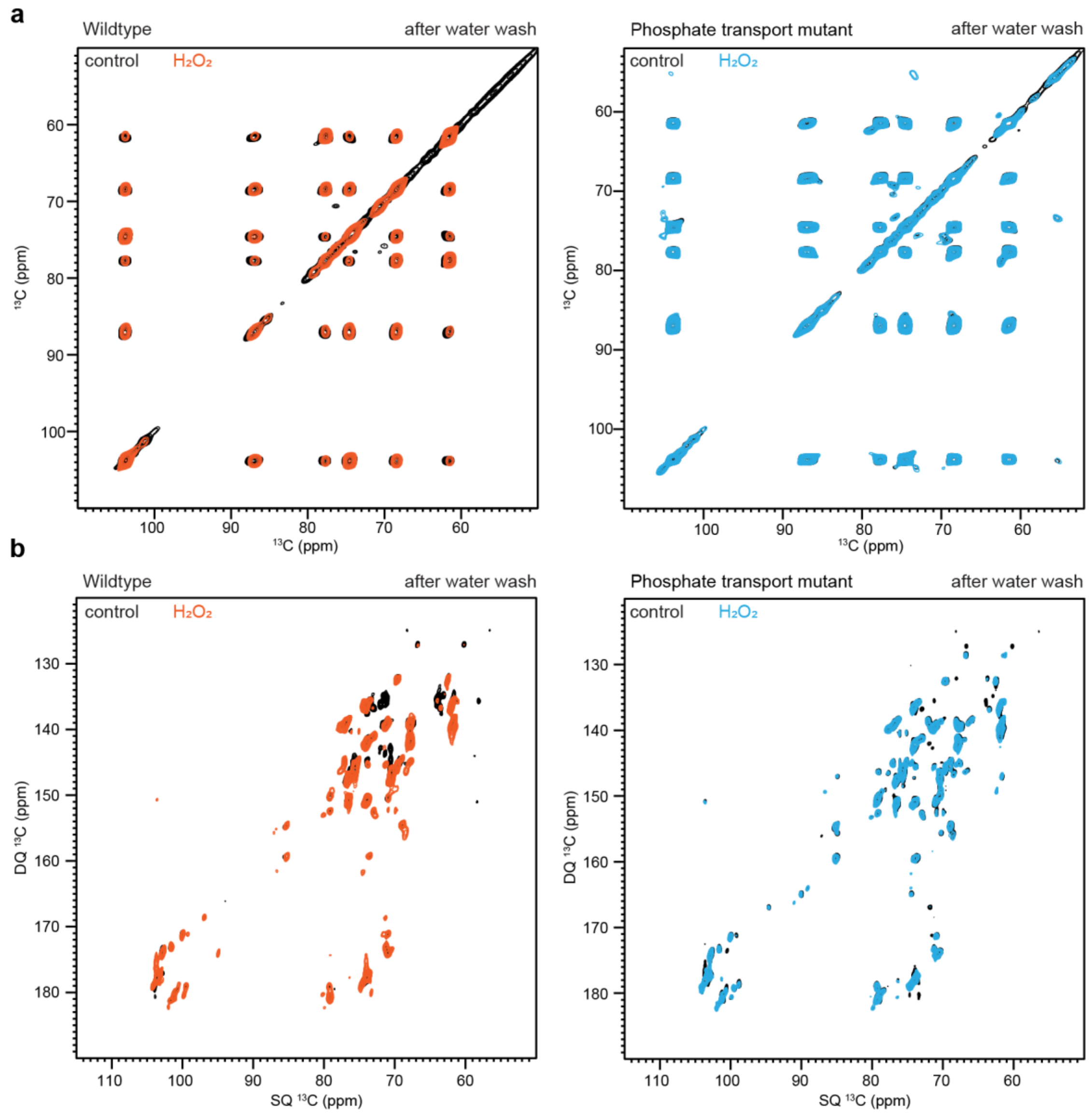


**Supplementary Figure 10. 2D $^{13}$C spectra of $H_2O_2$-treated *C. albicans* cell wall after washing.** (**a**) CP for selecting rigid polysaccharides. (**b**) DP spectra with a short recycle of 2s for selection of mobile components. (**c**) DP with long recycle delays (35s) for quantitative detection of all molecules. (**d**) Refocused INEPT experiments for probing the most dynamic molecules. The spectra show wildtype *C. albicans* control and $H_2O_2$ washed samples in black and orange, respectively and quadruple mutant control and $H_2O_2$ washed samples in black and cyan, respectively.

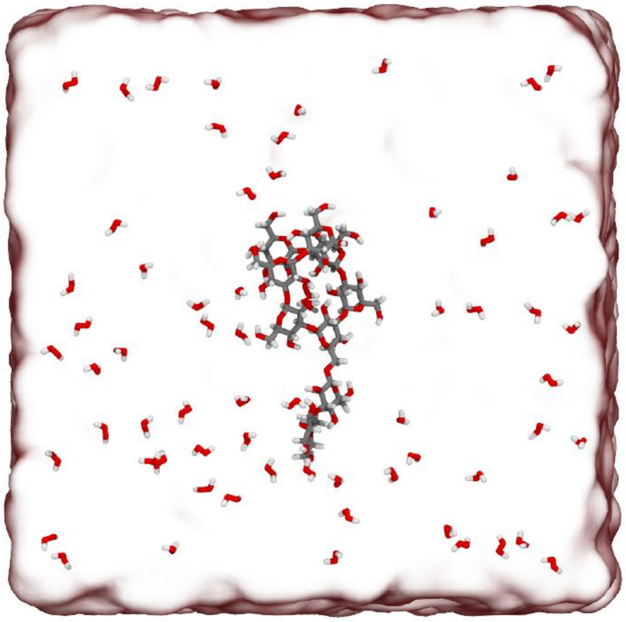

**Supplementary Figure 11**. **Molecular simulation snapshot.** A molecular system, featuring a small glycan chain (gray carbons, red oxygens, white hydrogens) within a water box (transparent surface) and a relatively high concentration of hydrogen peroxide molecules, shown here in a licorice representation.

**Supplementary Table 1. Solid-state NMR experiments and parameters.** To be quantitative, direct pulse (DP) experiments with 35 s long recycling delay were used. cross-polarization (CP), most rigid molecules. With DP and a shorter recycling delay of 2 seconds, suppress the rigid molecules from the spectra, and with Insensitive Nuclei Enhanced by Polarization Transfer (INEPT) the most mobile molecules were selected. For 2D $^{13}C$-$^{13}C$ correlation experiments allowed to resolve rigid intramolecular peaks. 2D DQ-SQ, DP J-INADEQAUTE and CP INADEQUATE spectra were used to detect through-bond correlations. The experimental parameters include the $^{1}H$ Larmor frequency, total experiment time (t), recycle delay (d1), number of scans (NS), The number of points for the direct (td2) and indirect (td1) dimensions, the acquisition time of the direct dimension (aq2) and the evolution time of indirect dimension (aq1), spectral width (sw1 and sw2), mixing time ($t_m$), increment delay (IN_F) and T filter times. * Indicates the water-polysaccharide spin diffusion. The processing parameters include the window function and associated parameters.

| | Acquisition parameters | | | | | | | | | | | | | Processing parameters | |
|---|---|---|---|---|---|---|---|---|---|---|---|---|---|---|---|
| Experiment | $\omega_{0,1H}$ (MHz) | t (h) | d1 (s) | NS | td2 | td1 | aq2 (ms) | aq1 (ms) | sw2 (ppm) | sw1 (ppm) | $t_m$ (ms) | IN_F (µs) | T filters | Window function | Parameter |
| 1D $^{13}C$ CP | 800 | 0.5 | 2.0 | 1024 | 3200 | | 16.0 | | 496.8 | | 1.0 | | | GM | LB-10, GB0.05 |
| 1D $^{13}C$ DP | 800 | 0.2 | 2.0 | 256 | 3200 | | 16.0 | | 496.8 | | | | | GM | LB-10, GB0.05 |
| 1D $^{13}C$ DP | 800 | 2.5 | 35.0 | 256 | 3200 | | 16.0 | | 496.8 | | | | | GM | LB-10, GB0.05 |
| 1D $^{13}C$ INEPT | 800 | 1.5 | 3.5 | 1024 | 3200 | | 16.0 | | 496.8 | | | | | GM | LB-10, GB0.05 |
| 1D $^{13}C$ T1 | 400 | 3.5 | 2.0 | 512 | 2000 | | 16.0 | | 623.3 | | | | $T_1(10^{-3}$-12 s) | GM | LB-10, GB0.05 |
| 1D $^{1}H$ T1ρ | 400 | 4.0 | 2.0 | 512 | 2000 | | 16.0 | | 623.3 | | | | SL ($10^{-3}$-19 ms) | GM | LB-10, GB0.05 |
| 2D $^{13}C$-$^{13}C$ CORD | 800 | 11.0 | 2.0 | 32 | 2800 | 600 | 14.0 | 7.5 | 496.8 | 198.7 | 53.0 | 25 | | QSINE | SSB 3.5 |
| 2D $^{13}C$-DP INADEQUATE | 800 | 6.0 | 2.0 | 16 | 2800 | 680 | 14.0 | 7.4 | 496.8 | 225.8 | | 22 | | | |
| 2D $^{13}C$-$^{13}C$ Water-edited | 400 | 8.0 | 2.0 | 64 | 2000 | 220 | 16.0 | 5.4 | 623.3 | 199.4 | $10^{-4}$/50* | 50 | $T_2$ $10^{-4}$ ms | QSINE | SSB 3.5 |

**Supplementary Table 2.** **$^{13}$C chemical shifts of biomolecules in *C. albicans* cell walls at ambient temperature.** Superscripts are used to denote different allomorphs. Not applicable (/). Unidentified (-). Branched (Br). Reducing end (O).

| Carbohydrate | | C1 | C2 | C3 | C4 | C5 | C6 | CO | $CH_3$ | N | Experiment | References |
|---|---|---|---|---|---|---|---|---|---|---|---|---|
| β-1,3-glucan | | 103.8 | 74.6 | 86.8 | 68.5 | 77.7 | 61.6 | / | / | / | $^{13}$C-$^{13}$C CORD | Shim *et al*. 2007[1] Fairweather *et al*. 2004[2] Saito *et al*. 1979[3] |
| Chitin | | 103.9 | 55.2 | 73.3 | 83.3 | 75.8 | 60.9 | 175.1 | 22.8 | / | | |
| β-1,3-6-glucan ($B^{Br}$) | | 103.2 | 73.9 | 85.4 | 69.1 | 75.8 | 69.2 | / | / | / | $^{13}$C DP J-INADEQUATE | Lowman et al. 2011[4] |
| β-1,6-glucan | | 103.7 | 74.2 | 76.4 | 70.5 | 75.2 | 69.5 | / | / | / | | |
| α-1,2-Mannan | a | 101.3 | 79.1 | 71.1 | 67.9 | 74.3 | 61.9 | / | / | / | | Latgé *et al*. 1994[5] Chakraborty *et al*. 2021[6] |
| | b | 99.2 | 79.4 | 71.8 | 67.7 | 73.8 | 62.2 | / | / | / | | |
| α-1,6-Mannan | | 102.8 | 70.8 | 73.7 | 67.5 | 71.2 | 66.8 | / | / | / | | |

**Supplementary Table 3. $^{13}C$-$T_1$ and $^{1}H$-$T_{1\rho}$ relaxation time constants with single exponential fitting.** A single exponential equation was used to fit the $T_1$ data $I(t) = e^{-t/T_I}$. A single exponential equation was used to fit the $T_{1\rho}$ data: $I(t) = e^{-t/T_{I\rho}}$. Error bars are standard deviations of the fit parameters.

| Sample Type | Cross peaks | $T_1$ (s) | Cross peaks | $T_{1\rho}$ (ms) |
|---|---|---|---|---|
| Wildtype | B1 | 1.4±0.07 | B1 | 10.4±0.61 |
| | B3 | 1.4±0.08 | B3 | 11.2±0.93 |
| | B5 | 1.2±0.08 | B5 | 11.3±0.79 |
| | B2 | 1.3±0.07 | B2 | 10.0±0.83 |
| | B4 | 1.2±0.06 | B4 | 11.5±1.01 |
| | Ch1 | 1.3±0.06 | Ch1 | 9.2±0.61 |
| | Ch4 | 2.1±0.27 | Ch4 | 5.3±1.08 |
| | Ch2 | 1.0±0.17 | Ch2 | 6.0±0.76 |
| | H3 | 1.1±0.09 | H3 | 6.9±0.79 |
| | H5 | 1.2±0.09 | H5 | 7.1±0.77 |
| | H6 | 1.2±0.12 | H6 | 7.0±0.94 |
| | Mn1 | 0.4±0.02 | Mn1 | 2.2±0.29 |
| | Mn2 | 1.3±0.10 | Mn2 | 4.8±0.80 |
| Phosphate transport mutant (JKC 2830) | B1 | 1.4±0.06 | B1 | 12.5±0.99 |
| | B3 | 1.4±0.03 | B3 | 13.6±1.24 |
| | B5 | 1.2±0.07 | B5 | 12.4±1.10 |
| | B2 | 1.4±0.10 | B2 | 11.2±0.94 |
| | B4 | 1.3±0.07 | B4 | 13.4±1.20 |
| | Ch1 | 1.4±0.06 | Ch1 | 12.6±0.88 |
| | Ch4 | 1.8±0.25 | Ch4 | 6.4±0.87 |
| | Ch2 | 1.8±0.23 | Ch2 | 9.8±1.21 |
| | H3 | 0.9±0.12 | H3 | 5.9±0.66 |
| | H5 | 1.5±0.18 | H5 | 8.1±0.93 |
| | H6 | 1.4±0.16 | H6 | 10.7±0.56 |
| | Mn1 | 0.6±0.05 | Mn1 | 4.8±0.32 |
| | Mn2 | 1.2±0.09 | Mn2 | 7.2±1.13 |

**Supplementary Table 4. $^{1}$H and $^{13}$C chemical shift of mobile carbohydrates of *C. albicans* wildtype cells.**

| Carbohydrates | forms | C1 | C2 | C3 | C4 | C5 | C6 | Reference |
|---|---|---|---|---|---|---|---|---|
| β-1,3-glucan (B) | a | 103.31<br>4.51 | 74.10<br>4.33 | 85.0<br>4.12 | 70.30<br>4.21 | 75.80<br>3.62 | 61.6<br>3.8, 3.9 | Shim *et al.* 2007[1]<br>Fairweather *et al.* 2009[7]<br>Saito *et al.* 1979[8] |
| | b | --- | 74.40 | 84.75<br>4.12 | 70.37 | --- | 61.79<br>3.80,3.90 | |
| | c | --- | 74.53<br>4.33 | 86.46<br>4.27 | 71.32<br>4.41 | --- | 62.35<br>3.83, 3.90 | |
| | d | --- | 74.2<br>4.32 | 85.95<br>4.21 | 71.38<br>4.42 | --- | 62.19<br>3.8, 3.9 | |
| β-1,3,6-glucan (Br) | a | 103.2<br>4.51 | 75.0<br>3.6 | 85.27<br>3.75 | 73.61<br>3.54 | --- | 69.13<br>3.92 | Lowman *et al.* 2011[9] |
| | b | | | 85.74<br>4.6 | | | 68<br>4.0 | |
| β-1,6-glucan (H) | | | | | | | | |
| | b | 103.54<br>4.52 | 74.01<br>3.33 | 76.3<br>3.5 | 69.9<br>3.47 | --- | --- | |
| | d | 102.63<br>4.46 | 74.4<br>3.34 | 76<br>3.5 | 69.56<br>3.85 | | | |
| α-1,6-Mannan ($Mn^{1,6}$) | | 102.75<br>5.15 | 70.92<br>4.08 | --- | 67.64<br>3.67 | 70.92<br>4.08 | --- | Latge *et al.* 1994[10]<br>Chakraborty *et al.* 2021[11]<br>Kuraoka *et al.* 2021[12] |
| α-1,2-Mannan ($Mn^{1,2}$) | a | 101.28<br>5.27 | 79.10<br>4.10 | 70.90<br>4.00 | 67.80<br>3.70 | 74.00<br>3.70 | 61.80<br>3.80/3.90 | |
| | b | 98.82<br>5.11 | 79.3<br>4.00 | 71.19<br>3.88 | 67.00<br>3.80 | --- | 61.98<br>3.84 | Kuraoka *et al.* 2021[12]<br>Kuraoka *et al.* 2018[13] |
| | c | 100.65<br>5.15 | 78.16<br>4.26 | 70.04<br>3.91 | 68.18<br>3.7 | 73.78<br>3.70 | 61.40<br>3.76, 3.9 | |
| | d | 101.2<br>5.38 | 79.00<br>4.10 | 70.80<br>4.10 | 67.96<br>3.70 | 73.77<br>3.70 | 61.76<br>3.76, 3.90 | |
| | e | 102.38<br>5.05 | 78.37<br>3.93 | 70.62<br>4.08 | 67.4<br>3.66 | 73.89<br>3.3 | 61.80<br>3.75, 3.9 | |

| | | | | | | | | |
|---|---|---|---|---|---|---|---|---|
| | f | 101.33<br>5.27 | 79.10<br>4.10 | 70.9<br>4.0 | 67.8<br>3.70 | 74.00<br>3.70 | 61.80<br>3.80,3.90 | |
| | g | 100.66<br>5.34 | 78.93<br>4.10 | 70.80<br>4.0 | 67.75<br>3.7 | 73.68<br>3.7 | 61.87<br>3.7, 3.9 | |
| | h | 102.57<br>5.03 | 78.76<br>3.93 | 70.69<br>4.08 | 66.3<br>3.90 | 73.95<br>3.7 | 61.80<br>3.75, 3.9 | |
| | i | 99.5<br>4.85 | 79.37<br>4.26 | --- | 67.88<br>3.62 | 72.43<br>3.69 | 61.51<br>3.78, 3.92 | |
| | j | 100.06<br>4.92 | --- | 71.1<br>3.87 | 67.2<br>3.61 | --- | 61.82<br>3.84<br>3.68 | |
| | k | 101.63<br>4.85 | 77.43<br>4.22 | 70.88<br>4.08 | 67.61<br>3.67 | 73.89<br>3.74 | 62.0<br>3.76, 3.9 | |
| | l | 101.71<br>4.93 | 77.87<br>4.25 | --- | 68.41<br>3.7 | 73.67<br>3.63 | 61.91<br>3.76, 3.9 | |
| | m | 94.79<br>5.54 | 78.77<br>4.19 | 69.66<br>4.0 | 67.56<br>3.67 | --- | 61.11<br>3.77, 3.83 | |
| Galactose/Glucose or their derivatives (Gl) | a | 90.00<br>5.90 | --- | 74.30<br>4.30 | 70.30<br>4.20 | -- | 61.70<br>3.80/ 3.90 | Fontaine *et al.* 2011[14] |
| | b | 96.84<br>5.46 | --- | 74.39<br>3.83 | 70.87<br>4.03 | 67.2<br>3.7 | 61.56<br>3.7, 3.83 | |
| | c | 94.61<br>4.90 | 71.60<br>3.94 | 73.65<br>3.66 | 67.00<br>3.60 | --- | 61.83<br>3.70, 3.80 | Archbald et al. 1981[15]<br>Fontaine *et al.* 2011[14] |
| | d | 94.95<br>5.18 | --- | 71.15<br>3.85 | 67.34<br>3.66 | --- | 61.7<br>3.70, 3.80 | |
| | e | 89.26<br>6.04 | --- | 74.32<br>4.33 | 71.24<br>4.42 | --- | 62.28<br>3.80, 3.90 | |
| | f | 88.51<br>5.87 | --- | 74.37<br>4.33 | 71.16<br>4.40 | --- | 62.46<br>3.83, 3.9 | |
| Glucose (Glc) | a (α) | 92.94<br>5.20 | 72.40<br>3.50 | 72.40<br>3.50 | 70.60<br>3.40 | --- | 61.56<br>3.70, 3.80 | Archbald *et al.* 1981[15] |
| | b (β) | 96.72<br>4.64 | 75.0<br>3.25 | 76.3<br>3.50 | 70.50<br>3.40 | --- | 61.66<br>3.70, 3.83 | |

**Supplementary Table 5. Experimental parameters used for both wildtype and quadruple mutant *C. albicans*.** Mobile region of the cell wall was characterized using scalar-coupling (*J*) based experiments on 800 MHz (18.8 T) spectrometer using 3.2 mm triple resonance MAS probe with the MAS frequency of 15 kHz.

<table>
<tr><th>Experiments</th><th>MAS</th><th>D1</th><th>NS</th><th>td2</th><th>td1</th><th>td3</th><th>aq2 (ms)</th><th>aq1 (ms)</th><th>aq3 (ms)</th><th>Decoupling/<br>Water suppression</th><th>J-evolution (ms)</th><th>DIPSI-3 (ms)</th><th>Expt. Time (h)</th></tr>
<tr><td></td><td></td><td></td><td></td><td></td><td></td><td></td><td></td><td></td><td></td><td rowspan="3">SPINAL-64<br>(rf 71.429 kHz)<br>WALTZ-16<br>(rf 17 kHz)<br>MISSISSIPI<br>(total duration)<br>40 ms<br>(rf 25.994 kHz</td><td></td><td></td><td></td></tr>
<tr><td>2D hCCH TOCSY (DIPSI-3)</td><td>15</td><td>2</td><td>32</td><td>2614<br>($^{1}$H)</td><td>512<br>($^{13}$C)</td><td>1</td><td>39.9</td><td>10.2</td><td>-</td><td rowspan="2">1.78 ($\tau_1$)<br>1.19 ($\tau_2$</td><td rowspan="2">25.5</td><td>9h42m</td></tr>
<tr><td>3D hCCH TOCSY (DIPSI-3)</td><td>15</td><td>2</td><td>8</td><td>2614<br>($^{1}$H)</td><td>128<br>($^{13}$C)</td><td>128<br>($^{13}$C)</td><td>39.9</td><td>2.56</td><td>2.56</td><td>77h53m</td></tr>
</table>

### Supplementary References


1 Shim, J. H. *et al.* Antitumor Effect of Soluble b-1, 3-Glucan from Agrobacterium sp. R259 KCTC 1019. *J. Microbiol. Biotechnol.* **17**, 1513-1520 (2007).

2 Fairweather, J. K., Him, J. L. K., Heux, L., Driguez, H. & Bulone, V. Structural characterization by $^{13}$C-NMR spectroscopy of products synthesized in vitro by polysaccharide synthases using $^{13}$C-enriched glycosyl donors: application to a UDP-glucose:(1→ 3)-b-d-glucan synthase from blackberry (Rubus fruticosus). *Glycobiology* **14**, 775-781 (2004).

3 Saitô, H., Ohki, T. & Sasaki, T. A 13C-nuclear magnetic resonance study of polysaccharide gels. Molecular architecture in the gels consisting of fungal, branched (1→ 3)-β-D-glucans (lentinan and schizophyllan) as manifested by conformational changes induced by sodium hydroxide. *Carbohydrate Research* **74**, 227-240 (1979).

4 Lowman, D. W. *et al.* New insights into the structure of (1→ 3, 1→ 6)-β-D-glucan side chains in the Candida glabrata cell wall. *PloS one* **6**, e27614 (2011).

5 Latge, J.-P. *et al.* Chemical and immunological characterization of the extracellular galactomannan of Aspergillus fumigatus. *Infection and immunity* **62**, 5424-5433 (1994).

6 Chakraborty, A. *et al.* A molecular vision of fungal cell wall organization by functional genomics and solid-state NMR. *Nat. Comm.* **12**, 6346 (2021).

7 Fairweather, J. K., Him, J. L. K., Heux, L., Driguez, H. & Bulone, V. Structural characterization by $^{13}$C-NMR spectroscopy of products synthesized in vitro by polysaccharide synthases using $^{13}$C-enriched glycosyl donors: application to a UDP-glucose:(1→ 3)-b-d-glucan synthase from blackberry (Rubus fruticosus). *Glycobiology* **14**, 775-781 (2004).

8 Saitô, H., Ohki, T. & Sasaki, T. A 13C-nuclear magnetic resonance study of polysaccharide gels. Molecular architecture in the gels consisting of fungal, branched (1→ 3)-b-D-glucans (lentinan and schizophyllan) as manifested by conformational changes induced by sodium hydroxide. *Carbohydr. Res.* **74**, 227-240 (1979).

9 Lowman, D. W. *et al.* New Insights into the Structure of (1→3,1→6)-β-D-Glucan Side Chains in the Candida glabrata Cell Wall. *PLoS One* **6**, e27614 (2011).

10 Latge, J. P. *et al.* Chemical and immunological characterization of the extracellular galactomannan of Aspergillus fumigatus. *Infect. Immun.* **62**, 5424-5433 (1994).

11 Chakraborty, A. *et al.* A molecular vision of fungal cell wall organization by functional genomics and solid-state NMR. *Nat. Commun.* **12**, 6346 (2021).

12 Kuraoka, T., Yamada, T., Takatsutsumi, Y., Ogawa, Y. & Kobayashi, H. Anomeric Proton and Carbon (H1-C1) NMR Chemical Shifts of Antigenic Mannans Obtained from Pathogenic Yeast Candida tropicalis. *Adv. Microbiol.* **11**, 296-301 (2021).

13 Kuraoka, T., Ishiyama, A., Oyamada, H., Ogawa, Y. & Kobayashi, H. Presence of O-glycosidically linked oligosaccharides in the cell wall mannan of Candida krusei purified with Benanomicin A. *FEBS Open Bio.* **9**, 129-136 (2018).

14 Fontaine, T. *et al.* Galactosaminogalactan, a New Immunosuppressive Polysaccharide of Aspergillus fumigatus. *PLoS Pathog.* **7**, e1002372 (2011).

15 Archbald, P. J., Fenn, M. D. & Roy, A. B. 13C-N.M.R. studies of D-glucose and D-galactose monosulphates. *Carbohydr. Res.* **93**, 177-190 (1981).
